# Amazonian biocultural heritage under climate change

**DOI:** 10.1101/2024.10.16.617268

**Authors:** Rodrigo Cámara-Leret, Patrick R. Roehrdanz, Jordi Bascompte

## Abstract

Amazonia harbors one fourth of the world’s plant diversity and over 300 Indigenous groups. So far, however, no study has assessed how climate change may simultaneously impact its biological and cultural heritage. To bridge this gap, we assembled a database on 5,833 utilized plant species and show that climate change will reduce more the ranges of utilized than of non-utilized species by 2070. Locally, Indigenous cultures may lose an average of 65% of their utilized plant species and 50% of their associated services from climate change. Regionally, the loss of threatened languages may result in a 41% reduction in the Amazonian knowledge pool. Overall, our results point to the strong climate vulnerability of Amazonian biocultural heritage.

---

Biologists call Amazonia the “Earth’s lungs”—for it harbors half of the world’s tropical forests. Anthropologists call it a “living library”—for the sophisticated knowledge of its Indigenous and local people. So far, however, no work has assessed how climate change may simultaneously impact Amazonia’s biological and cultural heritage. On the one hand, ecologists assessing the impacts of climate change have emphasized two scales: ecosystems (i.e., forest-savannah transitions (*1*)) and organisms (e.g., animals and plants (*2, 3*)). On the other hand, social scientists have addressed the importance of Indigenous and local knowledge for climate change mitigation (*4*), but not the fate of the species that matter to people. As a result of these separate investigations, our understanding of the climate change vulnerability of Amazonia’s unique biocultural heritage —of its plant species, plant services, and cultures— remains incipient (Fig. 1).

**Fig. 1.**
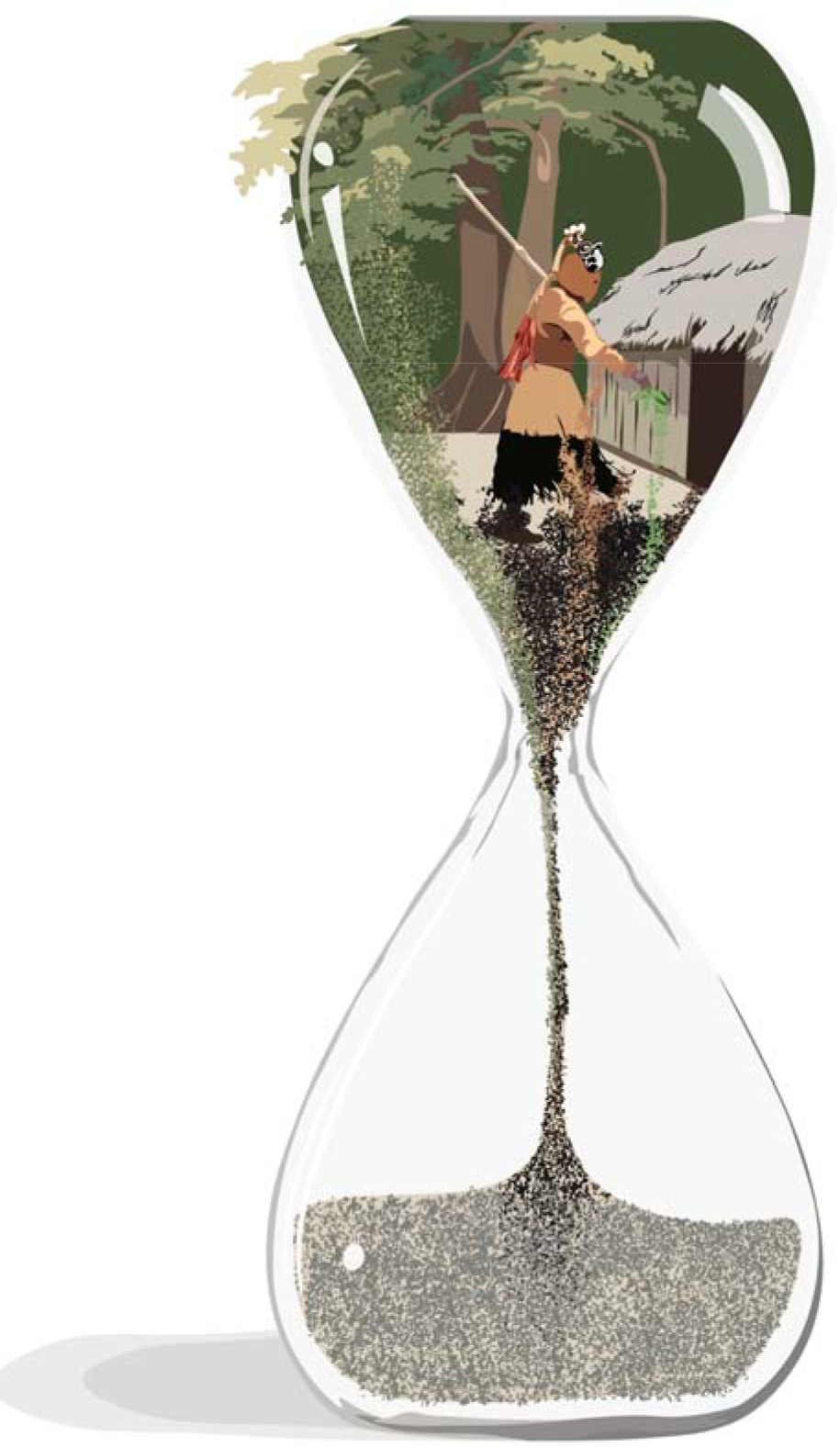
Linking biological and cultural heritage to study climate change impacts over time. The figure illustrates a sand clock for a particular culture in Amazonia, containing knowledge of plant species (biological heritage, green grains) and plant services (cultural heritage, black/brown grains). In this paper, we assess to what degree the biocultural heritage in this sand clock will be eroded by climate change. In our drawings of sand clocks in Fig. 3, this is quantified by the volume of sand grains occupying the lower globe.

To bridge this knowledge gap, and re-focus attention to those species that matter to people, we start by assessing to what degree climate change may impact the geographic range of utilized vs. non-utilized plant species. First, we reviewed 505 references on Amazonian utilized plants and recorded information for 5,833 utilized species from 74,602 literature reports. Next, we built species distribution models (SDMs) for 4,933 utilized species (85% of the utilized species in our database) and for 4,709 non-utilized species (46% of non-utilized species). We find two significant differences regarding their current distribution ranges. First, utilized plants have larger baseline ranges than non-utilized plants (t-Welch=28.14, p=1.09e-166) (fig. S1). Second, utilized plants have centroids that significantly differ in longitude (t-Welch=-12.67, p=1.70e-36) and latitude (t-Welch=-9.02, p=2.21e-19) (fig. S2 and S3). Projecting SDMs to eight climate scenarios by the year 2070, we then compared the mean change in the geographic range between utilized vs. non-utilized species. We find that climate change will differentially impact utilized plants. While non-utilized species are predicted to expand on average by 12%, utilized species are predicted to contract by a similar amount (Fig. 2).

**Fig. 2.**
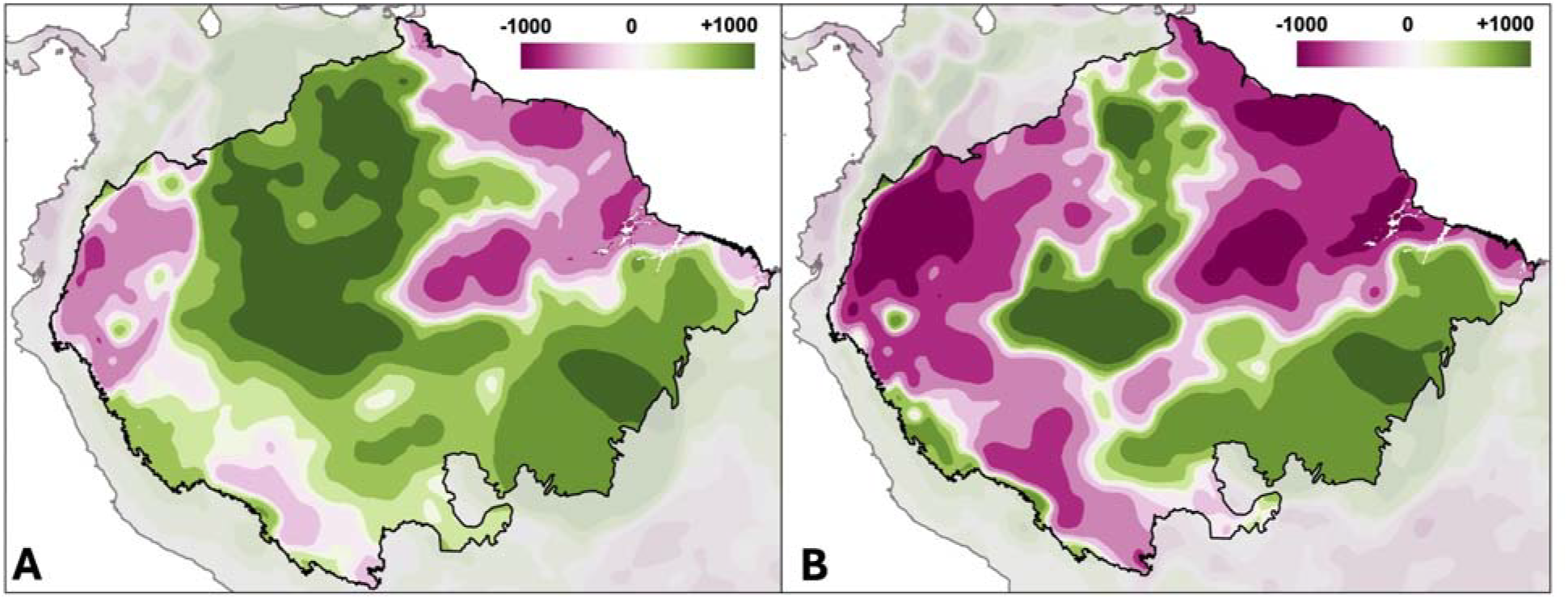
Climate change threatens biological heritage. Change in non-utilized (**A**) and utilized (**B**) plant species richness. In both panels, colors show the median change in species richness by 2070 under an extreme climate change scenario (RCP 8.5). The consequences of climate change will be more severe for the plant species used by people.

After having assessed the regional impact of climate change on utilized species, we now examine its effects at the local scale that matters to particular Indigenous cultures. Most cultures speak a distinct language and inhabit a particular territory —which linguists have mapped as language polygons (*5*)— allowing for local scale analyses. Because many languages are under-documented, here we only focus on 78 Indigenous languages with at least ten reported useful species in the literature (range: 10 to 1279 species; mean: 148; standard deviation: 224). For each language, we built an Indigenous knowledge network (*6*) to relate individual plant species (nodes in one set) to particular services (nodes on the other set) based on the knowledge (links) held by speakers of that language (see Materials and methods). Next, we quantified the climate change exposure of each Indigenous knowledge network by calculating the proportion of utilized species and services that may be driven locally extinct by 2070. We find that, on average, the proportion of plant species and services going locally extinct is 65% (range: 0 to 100%; sd: 22%) and 50% (range: 0 to 100%; sd: 1%), respectively (Fig. 3). The proportion of plant services lost will vary among cultures, even for cultures losing a similar fraction of species. Such loss is uncorrelated to the geographic longitude of languages, indicating that the biocultural impacts of climate change will be felt across the entire Amazon basin. The predicted climate change impacts are also decoupled from language threat, suggesting that when planning climate change mitigation actions in Amazonia, all cultures (not only the most endangered ones) need to be considered.

**Fig. 3.**
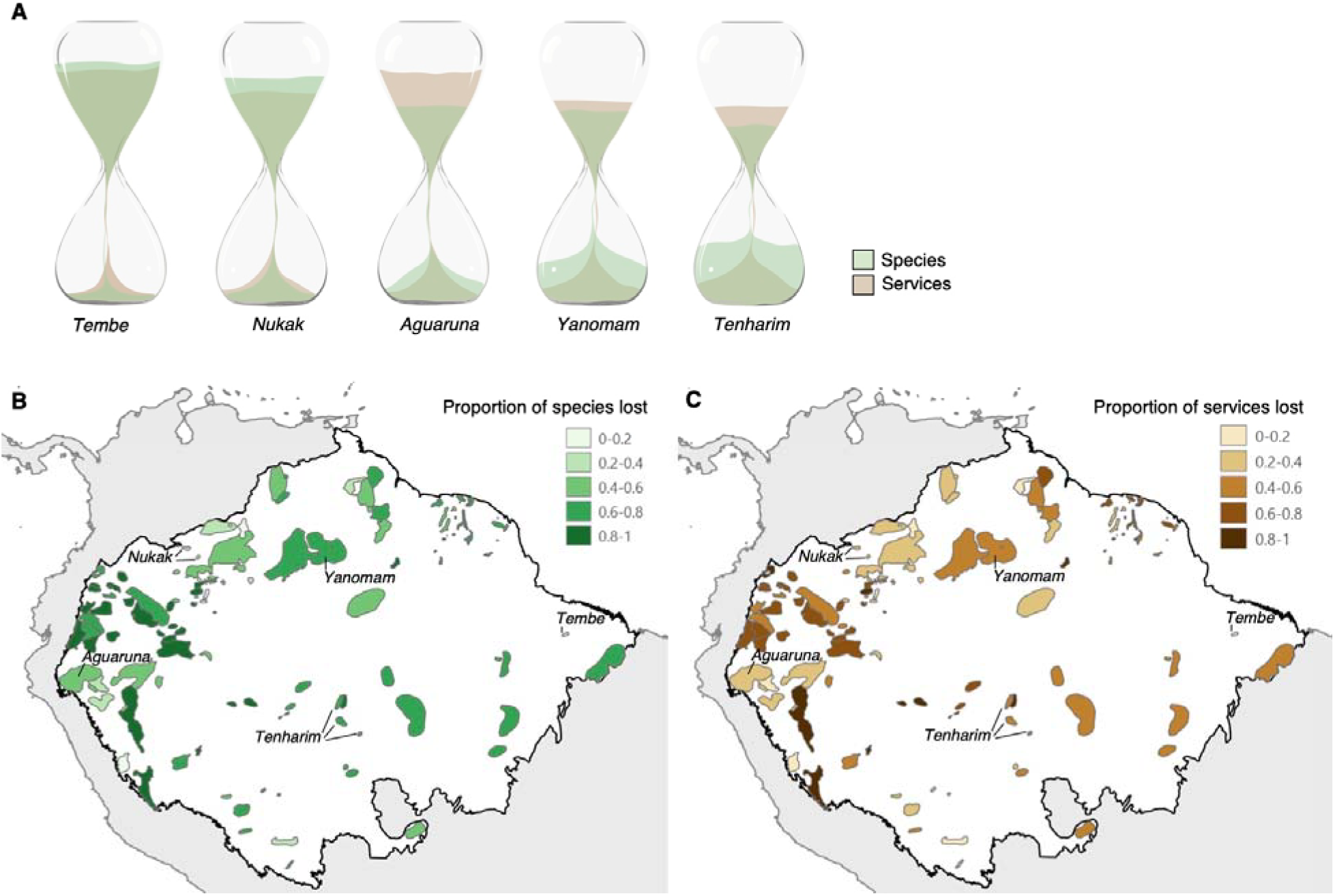
Climate change threatens cultural heritage. (**A**) Sand clocks show the predicted local extinction of plant services due to the local extinction of plant species from climate change. Five examples are shown to illustrate that local extinction of plant species and services varies across cultures. Proportion of plant species (**B**) or plant services (**C**) that are predicted to be lost within 78 Amazonian language polygons due to climate change by the year 2070.

So far, we have focused on the biocultural loss triggered by the extinction of plants, but the extinction of culture also matters. Indeed, cultural loss may undermine Indigenous knowledge networks at least as fast as biological loss (*6*). Specifically, language extinction may trigger the loss of medicinal knowledge, as most knowledge is unique to a single language (*7*). To assess how language extinction may impact Amazonian knowledge, we first build what we refer to as the indigenous knowledge metaweb (*6*) (i.e., the total knowledge about the services provided by all plant species in the study area). We then obtained linguistic threat data for all the languages in our sample from Ethnologue —the most comprehensive catalogue of the world’s languages (*5*)— and estimated the impact of language loss on the erosion of the Indigenous knowledge metaweb. From the 74,602 literature reports, 56% are from 138 Indigenous languages, of which 51% are threatened. By contrast, the other 44% are reports from non-specified languages whose threat status is unknown (81% of the 44%) or from non-Indigenous languages (e.g., Portuguese: 12%; Spanish: 7%). Accordingly, if threatened Indigenous languages vanish by the end of the century as predicted by linguists (*5*), the Amazonian metaweb would shrink by 44%. And yet, this estimate is conservative for two reasons. First, our sample includes nearly one third of Amazonian languages. Because 97% of non-sampled cultures speak threatened languages, including these in the analyses would further shrink the metaweb. Second, we (conservatively) classified all reports from non-specified languages as not threatened, but most Amazonian languages are, indeed, threatened. That is, if we would include non-specified language reports (assuming they are from threatened languages) the metaweb would shrink by 83%.

Here, we have shown that Amazonian peoples utilize at least 5,800 plant species, that utilized species may follow different climate change trajectories than non-utilized species, and that 60% of utilized species and 50% of their associated services may become locally extinct by 2070. Our findings are a conservative best-case scenario, as we do not consider land use change or mining, two major threats to Amazonia’s biocultural heritage (*8*). Hundreds of cultures and their languages have disappeared due to post-contact disease, epidemics, slavery and violence (*9*). As a result, their knowledge on the use and management of the world’s most biodiverse forest has been forever lost. Today, if languages continue to erode, much of the recorded knowledge will not be passed to the next generation, limiting the future wellbeing of the region’s inhabitants. Our results strongly hint to the possibility that the climate tipping point for Amazonia (*10*) will not only impact ecosystem and biological diversity, but also cascade across the biome’s astonishing and unique cultural heritage.

## Acknowledgments

We thank Martin Spinnler for helping locate books and articles, Lucia de la Torre for providing information from Ecuador, Inés Cámara Leret for the design of Fig. 1 and Fig. 3a, and members of the Cámara-Leret lab and Bascompte lab for feedback.

## Funding

Swiss National Science Foundation Starting Grant TMSGI3_211659 (RCL)

Swiss National Science Foundation project 310030_197201 (JB)

National Science Foundation BoCP-2225078 (PR)

John and Jody Arnhold University of California Santa Barbara - Conservation International Climate Solutions Collaborative (PR)

## Author contributions

Conceptualization: RCL, JB

Methodology: RCL, JB, PR

Investigation: RCL, JB, PR

Visualization: RCL, PR

Funding acquisition: RCL, JB

Writing – original draft: RCL, JB

Writing – review & editing: RCL, JB, PR

## References

1. B. M. Flores, M. Holmgren, C. Xu, E. H. Van Nes, C. C. Jakovac, R. C. Mesquita, M. Scheffer, Floodplains as an Achilles’ heel of Amazonian forest resilience. Proceedings of the National Academy of Sciences 114, 4442–4446 (2017).

2. L. P. Sales, M. Galetti, M. M. Pires, Climate and land-use change will lead to a faunal “savannization” on tropical rainforests. Global Change Biology 26, 7036–7044 (2020).

3. V. H. Gomes, I. C. Vieira, R. P. Salomão, H. ter Steege, Amazonian tree species threatened by deforestation and climate change. Nature Climate Change 9, 547–553 (2019).

4. V. Reyes-Garcia, D. Garcia-del-Amo, S. Álvarez-Fernández, P. Benyei, L. Calvet-Mir, A. B. Junqueira, V. Labeyrie, X. Li, S. Miñarro, V. Porcher, A. Porcuna-Ferrer, A. Schlingmann, C. Schunko, R. Soleymani, A. Tofighi-Niaki, M. Abazeri, E. M. N. A. N. Attoh, A. Ayanlade, J. Vieira Da Cunha Ávila, D. Babai, R. C. Bulamah, J. Campos-Silva, R. Carmona, J. Caviedes, R. Chakauya, M. Chambon, Z. Chen, F. Chengula, E. Conde, A. Cuní-Sanchez, C. Demichelis, E. Dudina, Á. Fernández-Llamazares, E. K. Galappaththi, C. Geffner-Fuenmayor, D. Gerkey, M. Glauser, E. Hirsch, T. Huanca, J. Tomás Ibarra, A. E. Izquierdo, L. Junsberg, M. Lanker, Y. López-Maldonado, J. Mariel, G. Mattalia, M. D. Miara, M. Torrents-Ticó, M. Salimi, A. Samakov, R. Seidler, V. Sharakhmatova, U. Babu Shrestha, A. Sharma, P. Singh, T. Ulambayar, R. Wu, I. S. Zakari, Indigenous peoples and local communities report ongoing and widespread climate change impacts on local social-ecological systems. Communications Earth & Environment 5, 29 (2024).

5. D. M. Eberhard, G. F. Simons, C. D. Fennig, Ethnologue: Languages of the World, 26th edn. Dallas, texas. TX: SIL International (2023).

6. R. Cámara-Leret, M. A. Fortuna, J. Bascompte, Indigenous knowledge networks in the face of global change. Proceedings of the National Academy of Sciences 116, 9913–9918 (2019).

7. R. Cámara-Leret, J. Bascompte, Language extinction triggers the loss of unique medicinal knowledge. Proceedings of the National Academy of Sciences 118, e2103683118 (2021).

8. D. M. Lapola, P. Pinho, J. Barlow, L. E. Aragão, E. Berenguer, R. Carmenta, H. M. Liddy, H. Seixas, C. V. Silva, C. H. Silva-Junior, A. A. C. Alencar, L. O. Anderson, D. Armenteras, V. Brovkin, K. Calders, J. Chambers, L. Chini, M. H. Costas, B. L. Faria, P. M. Fearnside, F. Ferreira, L. Gatti, V. H. Gutierrez-Velez, Z. Han, K. Hibbard, C. Koven, P. Lawrence, J. Pongratz, B. T. T. Portella, M. Rounsevell, A. C. Ruane, R. Schaldach, S. S. Da Silva, C. V. Randow, W. S. Walker, The drivers and impacts of Amazon forest degradation. Science 379, eabp8622 (2023).

9. M. J. Hamilton, R. S. Walker, D. C. Kesler, Crash and rebound of indigenous populations in lowland South America. Scientific Reports 4, 4541 (2014).

10. C. A. Nobre, L. D. S. Borma, “Tipping points” for the Amazon forest. Current Opinion in Environmental Sustainability 1, 28–36 (2009).

11. Rede Amazonica de Informacao Socioambiental Georreferenciada (2021), (available at https://www.amazoniasocioambiental.org/en/maps/).

12. J. Revilla, Plantas Úteis da Bacia Amazônica (SEBRAE-AM/INPA, 2002).

13. R. E. Schultes, R. F. Raffauf, The Healing Forest: Medicinal and Toxic Plants of the northwest Amazonia. (Dioscorides Press, 1990).

14. R. A. DeFilipps, S. L. Maina, J. Crepin, Medicinal Plants of the Guianas (Guyana, Surinam, French Guiana). (Department of Botany, National Museum of Natural History, Smithsonian, 2004).

15. L. De la Torre, H. Navarrete, P. Muriel, M. J. Maci a, H. Balslev, Enciclopedia de las Plantas Útiles del Ecuador (Herbario QCA de la Escuela de Ciencias Biológicas de la Pontificia Universidad Católica de Ecuador, 2008).

16. R. A. Rutter, Catálogo de Plantas Útiles de la Amazonia Peruana (Instituto Lingu□istico de verano, 1990).

17. P. Le Cointe, Árvores e Plantas Úteis: indígenas e aclimadas (Brasiliana, 1947).

18. A. B. Anderson, Os nomes e usos de palmeiras entre uma tribo de in dios Yanomama. Acta Amazónica 7, 5–13 (1977).

19. B. M. Boom, Ethnobotany of the Chácobo Indians, Beni, Bolivia. Advances in Economic Botany 4, 1–74 (1996).

20. G. Bourdy, S. DeWalt, L. C. De Michel, A. Roca, E. Deharo, V. Muñoz, L. Balderrama, C. Quenevo, A. Gimenez, Medicinal plants uses of the Tacana, an Amazonian Bolivian ethnic group. Journal of Ethnopharmacology 70, 87–109 (2000).

21. S. J. DeWalt, G. Bourdy, L.R. Chávez de Michel, C. Quenevo, Ethnobotany of the Tacana: Quantitative inventories of two permanent plots of northwestern Bolivia. Economic Botany 53, 237–260 (1999).

22. G. T. Prance, An ethnobotanical comparison of four tribes of Amazonian Indians. Acta Amazónica 2, 7–27 (1972).

23. M. Coelho-Ferreira, Medicinal knowledge and plant utilization in an Amazonian coastal community of Marudá, Pará state (Brazil). Journal of Ethnopharmacology 126, 159– 175 (2009).

24. I. B. da Costa, F. P. Bonfim, M. C. Pasa, D. A. Montero, Ethnobotanical survey of medicinal flora in the rural community Rio dos Couros, state of Mato Grosso, Brazil. Boleti n Latinoamericano y del Caribe de Plantas Medicinales y Aromáticas 16, 53–67 (2017).

25. L. L. C. Moraes, J. L. Freitas, J. R. Matos Filho, R. B. L. Silva, C. H. A. Borges, A. C. Santos, A ethno-knowledge of medicinal plants in a community in the eastern Amazon. Revista de Ciências Agrárias 42, 565–573 (2019).

26. P. Oliveira, B. Souza, Traditional knowledge of forest medicinal plants of Munduruku indigenous people-Ipaupixuna. European Journal of Medicinal Plants 31, 20–35 (2020).

27. G. X. de Paula Filho, A. F. Ribeiro, A. F. Moraes, W. F. Penha, W. L. Borges, R. H. S. Santos, Ethnobotanical knowledge on non-conventional food and medicinal plants in Rio Cajari Extractivist Reserve, Amazon, Brazil. Research Square, 10.21203/rs.3.rs-35316/v3 (2020).

28. K. A. Kainer, M. L. Duryea, Tapping women’s knowledge: Plant resource use in extractive reserves, Acre, Brazil. Economic Botany 46, 408–425 (1992).

29. C. T. Pedrollo, V. F. Kinupp, G. Shepard Jr, M. Heinrich, Medicinal plants at Rio Jauaperi, Brazilian Amazon: Ethnobotanical survey and environmental conservation. Journal of Ethnopharmacology 186, 111–124 (2016).

30. R. V. Ribeiro, I. G. C. Bieski, S. O. Balogun, D. T. de Oliveira Martins, Ethnobotanical study of medicinal plants used by ribeirinhos in the north Araguaia microregion, Mato Grosso, Brazil. Journal of Ethnopharmacology 205, 69–102 (2017).

31. C. La Rotta Cuéllar, Estudio etnobotánico sobre las especies utilizadas por la comunidad indi gena Miraña, Amazonas-Colombia (Bogotá: FEN-Colombia, 1986).

32. J. Sanz-Biset, J. Campos-de-la-Cruz, M.A. Epiquién-Rivera, S. Canigueral, A first survey on the medicinal plants of the Chazuta valley (Peruvian Amazon). Journal of Ethnopharmacology 122, 333–362 (2009).

33. W. Trujillo-C., M. Correa-Munera, Plants used by a Coreguaje indigenous community in the Colombian Amazon. Caldasia 32, 1–20 (2010).

34. C. Cerón, C. Reyes, P. Yépez, Mil y más plantas de la Amazonia ecuatoriana utilizadas por los Secoyas. Cinchonia 11, 13–205 (2011).

35. J. A. Echeverri, O. ‘Enokakuiodo’ Román-Jitdutjaaño, Witoto ash salts from the Amazon. Journal of Ethnopharmacology 138, 492–502 (2011).

36. D. Cárdenas, J. Arias, J. Vanegas, D. Jiménez, O. Vargas, L. Gómez, Plantas útiles y promisorias en la comunidad de Wacuraba (Caño Cuduyari) en el Departamento de Vaupés (Amazoni a colombiana). Bogotá, Instituto Amazónico de Investigaciones Cienti ficas-SINCHI (2007).

37. A. Lozano Balcázar, Los barbascos utilizados por los Ticuna del PNN Amacayacu, thesis, Uniandes (2005).

38. P. Giovannini, Medicinal plants of the Achuar (Jivaro) of Amazonian Ecuador: Ethnobotanical survey and comparison with other amazonian pharmacopoeias. Journal of Ethnopharmacology 164, 78–88 (2015).

39. C. Reyes, Conocimiento y uso de plantas en tres comunidades Kichwas: Yana Yaku, Loro Cachi y NinaAamarun, Pastaza-Ecuador. Cinchonia 15, 164–256 (2017).

40. M. N. Alexiades, Ethnobotany of the Ese Eja: Plants, health, and change in an Amazonian society (City University of New York, 1999).

41. V. Céline, P. Adriana, D. Eric, A. Joaquina, E. Yannick, L. F. Augusto, R. Rosario, G. Dionicia, S. Michel, C. Denis, Medicinal plants from the Yanesha (Peru): Evaluation of the leishmanicidal and antimalarial activity of selected extracts. Journal of Ethnopharmacology 123, 413–422 (2009).

42. G. Odonne, C. Valadeau, J. Alban-Castillo, D. Stien, M. Sauvain, G. Bourdy, Medical ethnobotany of the Chayahuita of the Paranapura basin (Peruvian Amazon). Journal of Ethnopharmacology 146, 127–153 (2013).

43. E. Fuentes, Los Yanomami y las plantas silvestres. Antropológica 54, 3–138 (1980).

44. B. M. Boom, Useful plants of the Panare indians of the Venezuelan Guayana. Advances in Economic Botany 8, 57–76 (1990).

45. P. B. Cavalcante, Frutas Comesti veis da Amazônia vol. I (Museu Paraense Emi lio Goeldi, 1972).

46. P. B. Cavalcante, Frutas Comesti veis da Amazônia vol. II (Museu Paraense Emi lio Goeldi, 1974).

47. P. B. Cavalcante, Frutas Comesti veis da Amazônia vol. III (Museu Paraense Emi lio Goeldi, 1974).

48. B. Krukoff, A. C. Smith, Rotenone-yielding plants of South America. American Journal of Botany 24, 573–587 (1937).

49. F. A. Flores, Notes on some medicinal and poisonous plants of Amazonian Peru. Advances in Economic Botany 1, 1–8 (1984).

50. M. Brandão, T. Grandi, E. Rocha, D. Sawyer, A. Krettli, Survey of medicinal plants used as antimalarials in the Amazon. Journal of Ethnopharmacology 36, 175–182 (1992).

51. V. Muñoz, M. Sauvain, G. Bourdy, J. Callapa, S. Bergeron, I. Rojas, J. Bravo, L. Balderrama, B. Ortiz, A. Gimenez, A search for natural bioactive compounds in Bolivia through a multidisciplinary approach: Part i. Evaluation of the antimalarial activity of plants used by the Chácobo Indians. Journal of Ethnopharmacology 69, 127–137 (2000).

52. S. Bertania, G. Bourdyb, I. Landaua, J. Robinsonc, P. Esterred, E. Deharo, Evaluation of French Guiana traditional antimalarial remedies. Journal of Ethnopharmacology 98, 45–54 (2005).

53. G. Odonne, F. Berger, D. Stien, P. Grenand, G. Bourdy, Treatment of leishmaniasis in the Oyapock basin (French Guiana): A K.A.P. survey and analysis of the evolution of phytotherapy knowledge amongst Wayãpi indians. Journal of Ethnopharmacology 137, 1228–1239 (2011).

54. B. Tomchinsky, L. C. Ming, V. F. Kinupp, A. de F. Hidalgo, F. C. M. Chaves, Ethnobotanical study of antimalarial plants in the middle region of the Negro River, Amazonas, Brazil. Acta Amazónica 47, 203–212 (2017).

55. W. Milliken, B. E. Walker, M.-J. R. Howes, F. Forest, E. N. Lughadha, Plants used traditionally as antimalarials in Latin America: Mining the tree of life for potential new medicines. Journal of Ethnopharmacology 279, 114221 (2021).

56. D. Cardoso, T. Särkinen, S. Alexander, A. M. Amorim, V. Bittrich, M. Celis, D. C. Daly, P. Fiaschi, V. A. Funk, L. L. Giacomin, R. Goldenberg, G. Heiden, J. Iganci, C. L. Kelloff, S. Knapp, H. Cavalcante de Lima, A. F. P. Machado, R. Manoel dos Santos, R. Mello-Silva, F. A. Michelangeli, J. Mitchell, P. Moonlight, P. L. Rodrigues de Moraes, S. A. Mori, T. SacramentoNunes, T. D. Pennington, J. Rubens Pirani, G. T. Prance, L. Paganucci de Queiroz, A. Rapini, R. Riina, C. A. Vargas Rincon, N. Roque, G. Shimizu, M. Sobral, J. R. Stehmann, W. D. Stevens, C. M. Taylor, M. Trovó, C. van den Berg, H. van der Werff, P. L. Viana, C. E. Zartman, R. C. Forzza, Amazon plant diversity revealed by a taxonomically verified species list. Proceedings of the National Academy of Sciences 114, 10695–10700 (2017).

57. H. Ter Steege, S. Mota de Oliveira, N. C. Pitman, D. Sabatier, A. Antonelli, J. E. Guevara Andino, G. A. Aymard, R. P. Salomão, Towards a dynamic list of Amazonian tree species. Scientific Reports 9, 1–5 (2019).

58. C. U. Ulloa, P. Acevedo-Rodriguez, S. Beck, M. J. Belgrano, R. Bernal, P. E. Berry, L. Brako, M. Celis, G. Davidse, R. C. Forzza, S. R. Gradstein, O. Hokche, B. León, S. León-Yánez, R. E. Magill, D. A. Neill, M. Nee, P. H. Raven, H. Stimmel, M. T. Strong, J. L. Villaseñor, J. L. Zarucchi, F. O. Zuloaga, P. M. Jørgensen, An integrated assessment of the vascular plant species of the Americas. Science 358, 1614–1617 (2017).

59. L. Hannah, P. R. Roehrdanz, P. A. Marquet, B. J. Enquist, G. Midgley, W. Foden, J. C. Lovett, R. T. Corlett, D. Corcoran, S. H. Butchart, B. Boyle, X. Feng, B. Maitner, J. Fajardo, B. J. McGill, C. Merow, N. Morueta-Holme, E. A. Newman, D. S. Park, N. Raes, J.-C. Svenning, 30% land conservation and climate action reduces tropical extinction risk by more than 50%. Ecography 43, 943–953 (2020).

60. S. J. Phillips, R. P. Anderson, R. E. Schapire, Maximum entropy modeling of species geographic distributions. Ecological Modelling 190, 231–259 (2006).

61. X. Feng, C. Merow, Z. Liu, D. S. Park, P. R. Roehrdanz, B. Maitner, E. A. Newman, B. L. Boyle, A. Lien, J. R. Burger, M. M. Pires, P. M. Brando, M. B. Bush, C. N. H. McMichael, D. M. Neves, E. I. Nikolopoulos, S. R. Saleska, L. Hannah, D. D. Breshears, T. P. Evans, J. R. Soto, K. C. Ernst, B. J. Enquist, How deregulation, drought and increasing fire impact Amazonian biodiversity. Nature 597, 516–521 (2021).

62. C. Merow, M. J. Smith, T. C. Edwards Jr, A. Guisan, S. M. McMahon, S. Normand, W. Thuiller, R. O. Wu est, N. E. Zimmermann, J. Elith, What do we gain from simplicity versus complexity in species distribution models? Ecography 37, 1267–1281 (2014).

63. C. Merow, M. J. Smith, J. A. Silander Jr, A practical guide to MaxEnt for modeling species’ distributions: What it does, and why inputs and settings matter. Ecography 36, 1058–1069 (2013).

64. J. M. Kass, R. Muscarella, P. J. Galante, C. L. Bohl, G. E. Pinilla-Buitrago, R. A. Boria, M. Soley-Guardia, R. P. Anderson, ENMeval 2.0: Redesigned for customizable and reproducible modeling of species’ niches and distributions. Methods in Ecology and Evolution 12, 1602– 1608 (2021).

65. R. J. Hijmans, S. E. Cameron, J. L. Parra, P. G. Jones, A. Jarvis, Very high resolution interpolated climate surfaces for global land areas. International Journal of Climatology: A Journal of the Royal Meteorological Society 25, 1965–1978 (2005).

66. A. Trabucco, R. J. Zomer, Global aridity index and potential evapotranspiration (ET0) climate database v2. CGIAR Consort Spat Inf. 10, m9 (2018).

67. T. Hengl, J. Mendes de Jesus, G. B. Heuvelink, M. Ruiperez Gonzalez, M. Kilibarda, A. Blagotić, W. Shangguan, M. N. Wright, X. Geng, B. Bauer-Marschallinger, M. A. Guevara, R. Vargas, R. A. MacMillan, N. H. Batjes, J. G. B. Leenaars, E. Ribeiro, I. Wheeler, S. Mantel, B. Kempen, SoilGrids250m: Global gridded soil information based on machine learning. PLoS One 12, e0169748 (2017).

68. R. T. Corlett, K. W. Tomlinson, Climate change and edaphic specialists: Irresistible force meets immovable object? Trends in Ecology & Evolution 35, 367–376 (2020).

69. B. S. Maitner, B. Boyle, N. Casler, R. Condit, J. Donoghue, S. M. Durán, D. Guaderrama, C. E. Hinchliff, P. M. Jørgensen, N. J. Kraft, B. McGill, C. Merow, N. Morueta-Holme, R. K. Peet, B. Sandel, M. Schildhauer, S. A. Smith, J.-C. Svenning, B. Thiers, C. Violle, S. Wiser, B. J. Enquist, The bien r package: A tool to access the botanical information and ecology network (BIEN) database. Methods in Ecology and Evolution 9, 373–379 (2018).

70. B. Enquist, B. Boyle, SALVIAS—the SALVIAS vegetation inventory database. Biodiversity and Ecology 4, 288 (2012).

71. B. J. Enquist, R. Condit, R. K. Peet, M. Schildhauer, B. M. Thiers, “Cyberinfrastructure for an integrated botanical information network to investigate the ecological impacts of global climate change on plant biodiversity” (PeerJ Preprints, 2016): https://peerj.com/preprints/2615/

72. E. Fegraus, Tropical Ecology Assessment and Monitoring network (TEAM network). Biodiversity and Ecology 4, 287 (2012).

73. R. K. Peet, M. T. Lee, M. D. Jennings, D. Faber-Langendoen, VegBank–a permanent, open-access archive for vegetation-plot data. Biodiversity and Ecology 4, 233–241 (2012).

74. K. J. Anderson-Teixeira, S. J. Davies, A. C. Bennett, E. B. Gonzalez-Akre, H. C. Muller-Landau, S. Joseph Wright, K. Abu Salim, A. M. Almeyda Zambrano, A. Alonso, J. L. Baltzer, Y. Basset, N. A. Bourg, E. N. Broadbent, W. Y. Brockelman, S. Bunyavejchewin, D. F. R. P. Burslem, N. Butt, M. Cao, D. Cardenas, G. B. Chuyong, K. Clay, S. Cordell, H. S. Dattaraja, X. Deng, M. Detto, X. Du, A. Duque, D. L. Erikson, C. E.N. Ewango, G. A. Fischer, C. Fletcher, R. B. Foster, C. P. Giardina, G. S. Gilbert, N. Gunatilleke, S. Gunatilleke, Z. Hao, W. W. Hargrove, T. B. Hart, B. C.H. Hau, F. He, F. M. Hoffman, R. W. Howe, S. P. Hubbell, F. M. Inman-Narahari, P. A. Jansen, M. Jiang, D. J. Johnson, M. Kanzaki, A. Rahman Kassim, D. Kenfack, S. Kibet, M. F. Kinnaird, L. Korte, K. Kral, J. Kumar, A. J. Larson, Y. Li, X. Li, S. Liu, S. K.Y. Lum, J. A. Lutz, K. Ma, D. M. Maddalena, J.-R. Makana, Y. Malhi, T. Marthews, R. Mat Serudin, S. M. McMahon, W. J. McShea, H. R. Memiaghe, X. Mi, T. Mizuno, M. Morecroft, J. A. Myers, V. Novotny, A. A. de Oliveira, P. S. Ong, D. A. Orwig, R. Ostertag, J. den Ouden, G. G. Parker, R. P. Phillips, L. Sack, M. N. Sainge, W. Sang, K. Sri-ngernyuang, R. Sukumar, I-F. Sun, W. Sungpalee, H. Sathyanarayana Suresh, S. Tan, S. C. Thomas, D. W. Thomas, J. Thompson, B. L. Turner, M. Uriarte, R. Valencia, M. I. Vallejo, A. Vicentini, T. Vrška, X. Wang, X. Wang, G. Weiblen, A. Wolf, H. Xu, S. Yap, J. Zimmerman, CTFS-forest GEO: A worldwide network monitoring forests in an era of global change. Global Change Biology 21, 528–549 (2015).

75. G. Csardi, T. Nepusz, The igraph software package for complex network research. Complex Systems, 1695 (2006).

