## Supplementary Material for "Amazonian biocultural heritage under climate change"

**The PDF file includes:**

Materials and Methods

Figs. S1 to S3

Tables S1

Materials and Methods

Study area

We delimit our study area of the Amazon basin following the biogeographic limits proposed by the Amazon Network of Georeferenced Socio-Environmental Information - RAISG (*11*).

Plants used by Amazonian cultures

To understand how climate change may affect Amazonia’s biocultural heritage, we first compiled bibliographic information on the plants that Indigenous and local communities utilize (table S1). We gathered four different types of information: i) Regional compilations for the Amazon basin (*12*), northwest Amazonia (*13*), and the Guianas (*14*), ii) country-level compilations, which exist for Ecuador (*15*), Peru (*16*), and Brazil (*17*), iii) Individual Indigenous groups (e.g., Refs. *18–44*), and iv) Specific plant services, e.g., Food (e.g., Refs. *45–47*) and Medicine (e.g., Refs. *48–55*). Our review represents the most comprehensive ethnobotanical assessment in Amazonia, covering one third of its Indigenous cultures. As more field ethnobotanical studies are conducted, the number of studied cultures will continue to increase, as will the number of utilized species (since many Amazonian Indigenous groups in the literature are superficially studied). To address the latter, we focused our local-scale analyses on only the best-studied cultures (see below).

To unify nomenclature and verify the geographic origin of species, we used the checklists to the vascular plants of Amazonia (*56*), Amazonian trees (*57*), Vascular Plants of the Americas (*58*), and Plants of the World Online (http://www.plantsoftheworldonline.org). All nonnative species or genus-level reports were discarded. We verified Indigenous group names and recorded their associated language and language threat status using Ethnologue (*5*).

Our database contains 74,602 use reports from 5,833 utilized native plant species, 1,291 genera, and 189 plant families. We defined a “use report” as the concatenation of a plant species with plant part, use category, and use subcategory (e.g., *Qualea homosepala* + Bark + Medicine + Blood and cardiovascular system). Of the use categories analyzed, Medicine contained 44% of all use reports, followed by Human Food (16%) and Construction (11%). The surveyed literature had information for 138 Indigenous languages that we could classify into 18 language families and seven language isolates. Five languages had 47% of all use reports (Kichwa (17%), Waorani (11%), Secoya (8%), Cofan (7%), and Shuar (4%)), whereas five language families account for 58% of the languages in our sample (Arawak, Tupi, Cariban, Pano-Tacana, and Tucano).

Species range models

Modelled species ranges were selected from the dataset used in Hannah et al. 2020 (*59*). Briefly, species distribution models were produced with Maxent (*60*) for species with >10 unique occurrence that were also on the list of Amazonian useful plants. Additionally, modelled plant species with >20% of the total modeled range in the Amazon basin were selected as the control group (*61*). The final number of species used in this analysis was 4,933 Amazonian utilized plant species and 4,709 control group species – for a combined total of 9,642 species. Maxent settings followed the recommendations of Merow et al. (*62*, *63*) to produce relatively less complex models (e.g., limiting features to linear, quadratic, and product functions) to minimize overfitting. Modelling domains were limited to a spatial buffer within 500 km of any valid occurrence record. This likewise limited the future projected ranges to within 500 km of any verified observation. Background sampling was a random sample of 10,000 points within the buffered modelling domain. Five model replicates were used in fitting the model and an average of the five replicates was used for the final species model. Parameters from the final model were used to project species suitability for both baseline and future climate scenarios. Thirty percent of occurrence records were randomly reserved to assess model performance. Model performance was assessed through area under the receiver operating curve (AUC) using the ENMeval R package (*64*). Binary range maps were produced from continuous model outputs using maximum sensitivity + specificity logistic threshold.

Environmental variables

We chose the following bioclimatic variables from downscaled (2.5 arc-minute resolution) climate normals available in Worldclim ver. 1.4 database (*65*) for both baseline (1960–1990) and future (2060–2080) climates: mean annual temperature, mean diurnal temperature range, seasonality of temperature, minimum temperature of the coldest month, mean annual precipitation, and seasonality of precipitation. We also used an accumulated aridity index that is the sum of the monthly aridity (annual precipitation - PET) for the maximum run of consecutive months where PET>precipitation. The accumulated aridity index was derived from global monthly extra-terrestrial solar radiation data from Ref. (*66*) and monthly maximum temperature, minimum temperature and precipitation from Worldclim ver. 1.4 (*65*). All future environmental variables were obtained for RCP8.5 (representing a high climate forcing scenario). Results presented here are for eight global climate models available that were a priori selected for their performance across a pan-tropical modeling domain. Soil variables used in species distribution models were depth to bedrock, pH, clay proportion, silt proportion and bulk density. All soil-related variables were obtained from Soilgrids ver. 1.0 (*67*). Variables with multiple strata available are the mean of the top 1m (strata 1–4). Soil variables were included as it has been shown that climate change analyses that do not incorporate soil variability can misrepresent edaphic specialists (*68*).

Plant Occurrence Records

Vascular plant data was extracted from the BIEN ver. 4.1 database using the RBIEN package (*69*). BIEN employs taxonomic name resolution, geographic centroid detection, geographic name resolution, and native species resolution services as a means of cleaning and standardizing occurrence records. Records retained for modelling were determined to: 1) be within a species native range; 2) not represent a centroid of a political boundary; 3) not be flagged as cultivated or introduced. The BIEN data mainly comprise herbarium collections, ecological plots and surveys (*70–74*). For details of specimen data sources see Maitner et al. (*69*). A full listing of the herbaria data used is given in Supplementary Methods. The observations in the BIEN database are the product of contributions by 1,076 different data contributors, including numerous individual herbaria, and data indexers of herbarium or plot data. Of the herbaria, 550+ are listed in Index Herbariorum. Additionally, BIEN 4.1 included data from RAINBIO, TEAM, The Royal Botanical Garden of Sydney, Australia, and NeoTropTree. Plot data within BIEN are from the CVS, NVS, SALVIAS, VEGBANK, CTFS, FIA, MADIDI, and TEAM data networks and datasets. Please refer to the BIEN database documentation for further details.

Climate change and language extinction impact on Indigenous knowledge networks

To assess how climate change may impact utilized vs. non-utilized plant species, we first built SDMs for 4,933 utilized and 4,709 non-utilized species, respectively (see Species range models above). Next, we compared changes in the average range of utilized vs. non-utilized species between the present and 2070 according to eight climate scenarios. To understand the impacts of language extinction across Amazonia, we first built an Indigenous knowledge metaweb which contains the total pool of knowledge in our database. The metaweb included 5,833 species, 709 plant services, and 36,152 interactions. We then obtained language threat data for all languages in our sample from Ethnologue (see above). This information was then used to calculate what proportion of the metaweb will remain after all threatened languages are lost. That is, when threatened languages vanish, the metaweb only contains knowledge associated to non-Indigenous groups who speak Portuguese and Spanish and to non-specified Indigenous languages which are assumed to be non-threatened. However, since this assumption is conservative, we repeated these analyses by classifying all non-specified languages as threatened. Finally, to understand how climate change may affect individual cultures, we built Indigenous knowledge networks (*6*) for the best-studied 78 languages which had at least 10 species reports in the literature. We modeled changes in utilized species in each network by comparing how many of the current useful species persist in the future within the Ethnologue language polygons associated to each network. This information was then used to calculate how the localized loss of species (and associated services) will affect network connectance by 2070 using the igraph R package (*75*).

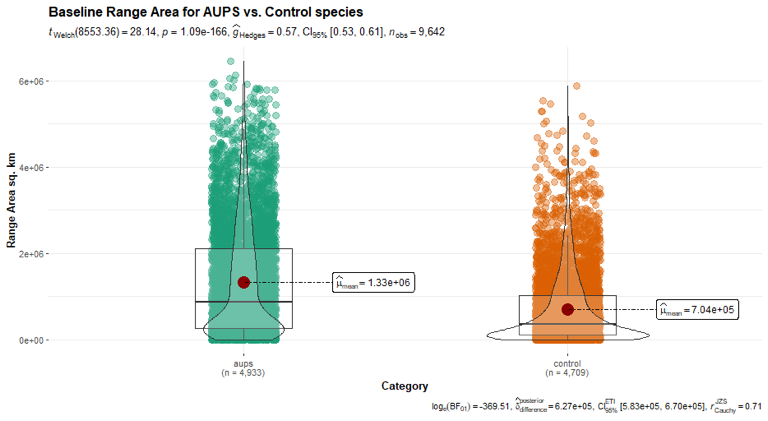

Fig. S1. Utilized plants have larger baseline ranges (left) than non-utilized plants (right).

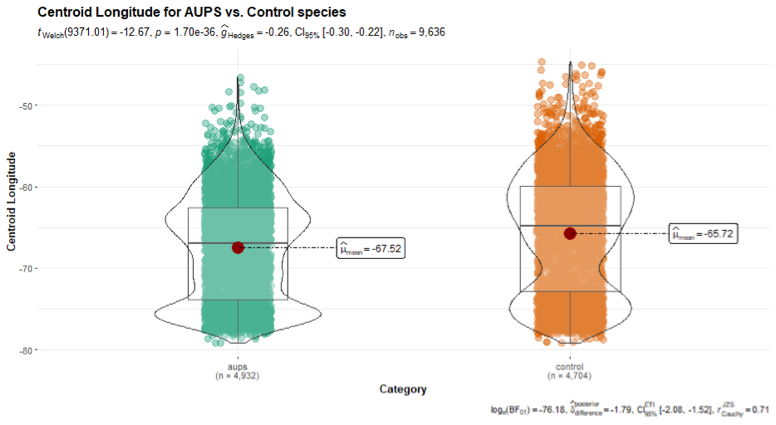

Fig. S2. Utilized plants (left) have centroids that significantly differ in longitude than non-utilized plants (right).

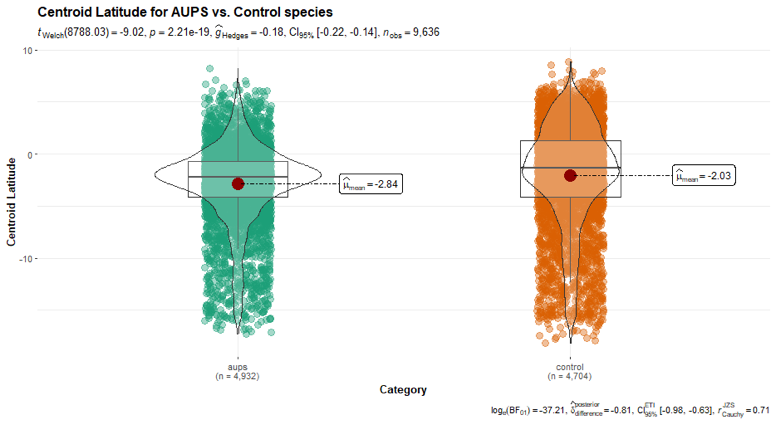

Fig. S3. Utilized plants (left) have centroids that significantly differ in latitude than non-utilized plants (right).

Table S1. References reviewed in this paper.

**Reference**

ACEER, cited as source in: Duke, J.A., & Wain, K.K. (1981) Medicinal Plants of the World. Computer index with more than 85,000 entries, 3 vols.

Acero-Duarte, L.E. (1979) Principales plantas útiles de la Amazonia colombiana. Editora Guadalupe, Bogotá

Acevedo-Rodriguez, P. (1990) The Occurrence of Piscicides and Stupefactants in the Plant Kingdom. Advances in Economic Botany vol. 8: 1-23

Acosta-Solis, M. (1952) Las fibras y lanas vegetales en el Ecuador. Contribución No. 21, Instituto Ecuatoriano de Ciencias Naturales. Casa de la Cultura Ecuatoriana, Quito

Aguilar, Z. (2006) Influence of the Huaorani on the conservation of Oenocarpus bataua, Arecaceae in Yasuni National Park and Biosphere Reserve, Amazonian Ecuador. Lyonia 10: 83-90

Aguirre, G. (2006) Plantas medicinales utilizadas por los indígenas Mosetén-Tsimane´ de la comunidad Asunción del Quiquibey, en la RB-TCO Pilón Lajas, Beni, Bolivia. Facultad de Ciencias Puras y Naturales. Universidad Mayor de San Andres

Ahlbrinck, W. (1931). Encyclopaedie der Kari√ïben, Amsterdam, 1931; traduction franpaise de Doude van Herwijnen, Institut Gdographique National, Paris, 1956.

Alarcón, R. (1988) Etnobotánica de los Quichuas de la Amazonia ecuatoriana. Miscelánea Antropológica Ecuatoriana, Serie Monográfica (Ecuador) 7

Alarcon, E.D.B. (2013). Estudio etnobotánico y etnofarmacológico de las especies aromáticas usadas en ceremonias de ayahuasca por la etnia Huarayo (Puerto Maldonado). Tesis. Universidad Nacional de San Antonio Abad del Cusco, Peru.

Alarcon, R. (1994) El taller "Etnobotánica y valoración económica de los recursos florísticos silvestres". Etnobotánica, Valoración Económica y Comercialización de Recursos Florísticos Silvestres en el Alto Napo, Ecuador, Ecociencia, Quito

Albán, J. (1994) La mujer y las plantas útiles silvestres en la comunidad Cocama-Cocamilla de los ríos Samiria y Marañon.. Informe de Proyecto. Wild World Life Foundation-Biodiversity Support Program 7560, Lima

Albarracin, J. & Caycedo, A.O. (2001). La pesca entre los ticuna: história, técnicas y ecossistemas. Revista IMANI MUNDO: estúdios em la Amazônia colombiana. Universidade Nacional de Colombia, Letícia.

Alexiades, M. N. (1999). Ethnobotany of the Ese Eja: Plants, health, and change in an Amazonian society (Doctoral dissertation, City University of New York).

Allen, P.H. (1947) Indians of Southeastern Colombia. Geographical Review 37: 567-582

Alves de Souza, J.M. & Chau Ming, L. (2016) Plantas medicinais utilizadas por seringueiros na Reserva Extrativista São Luiz do Remanso, Acre. Chapter 13. In: da Silva Ramos, J. I., Cavalcante, O. P., da Silva, J. O., de Oliveira, M. A., Fernandes, C. C., da Silva, J. P., & Uchôa, J. M. S. (2016). Etnobotânica e botânica econômica do Acre ISBN: 978-85-8236-027-9 Copyright© Edufac 2016, Amauri Siviero, Lin Chau Ming, Marcos Silveira, Douglas Daly, Richard Wallace (Organizadores) Editora da Universidade Federal do Acre-Edufac.

Anderson, A. B. (1977). Os nomes e usos de palmeiras entre uma tribo de índios Yanomama. Acta Amazonica, 7(1), 5-13.

Anderson, P.J. (2004) The social context for harvesting Iriartea deltoidea (Arecaceae). Economic Botany 58: 410-419

Andoque, H., Andoque, D., Andoque, M., & Andoque, H. (2009). Plantas medicinales de la Gente de Hacha. Universidad Nacional de Colombia-Sede Amazonia.

Andrade, J. N., Costa Neto, E. M., & Branda_o, H. (2015). Using ichthyotoxic plants as bioinsecticide: A literature review. Revista Brasileira de Plantas Medicinais, 17, 649-656.

Antezana, L. (1976) Palmeras nativas de Bolivia de valor económico. Simposio internacional sobre plantas de interés económico de la flora amazónica, Turrialba, Costa Rica

Antolinez, L. D. (1999). La alimentación en la Amazonía: Estudio de caso entre los coreguajes. Graduate thesis. Departamento de Filología e Idiomas, Facultad de Ciencias Humanas. Universidad Nacional de Colombia, Bogotá.

Araujo-Murakami, A. (2019). Barbascos y Curare en Bolivia. Kempffiana 15 (1): 57-63

Armesilla, P.J. (2006) Usos de las palmeras (Arecaceae),en la Reserva de la Biosfera-Tierra Comunitaria de Orígen Pilón Lajas, (Bolivia). Facultad de Ciencias. Universidad Autónoma de Madrid, Madrid.

Arnaud, E. (1975) Os índios Gaviões de Oeste Pacificação e integração . Publicações avulsas do Museu Goeldi (28) : 1-86. Pg. 27

Arrais, F. D. C. L., Viana, R. H. O., Silva, W. A., Pereira, B. L., & Ferreira, G. (2017). Levantamento etnobotânico nas margens do córrego Machado-Palmas, Tocantins, Brazil. FLOVET-Boletim do Grupo de Pesquisa da Flora, Vegetação e Etnobotânica, 1(9).

Aublet, F. (1775). Histoire de la Guayana Française. Paris, Pierre-Francois Didot jeune, Libraire de la Faculté de Médicine. 2 : 871

Ayala, F. (1984). Notes on Some Medicinal and Poisonous Plants of Amazonian Peru. Advances in Economic Botany 1: 1-8.

Báez, S. (1998) Dictionary of plants used by the Canelos-Quichua. People and biodiversity- Two case studies from the Andean foothills of Ecuador. Centre for research on cultural and biological diversity of Andean rainforests (DIVA). Rønde

Báez, S., and Å. Backevall (1998) Dictionary of plants used by the Shuar of Makuma and Mutints. People and biodiversity- Two case studies from the Andean foothills of Ecuador. Centre for research on cultural and biological diversity of Andean rainforests (DIVA). Rønde

Balée, W. (1987). A etnobotânica quantitativa dos índios Tembé (Rio Gurupi, Pará). Boletim do Museu Paraense Emilio Goeldi. Nova serie Botánica, 3(1), 29-50.

Balée, W. (1989). The culture of Amazonian forests. Advances in economic botany, 1-21.

Balée, W. L. (1994). Footprints of the forest: Ka'apor ethnobotany-the historical ecology of plant utilization by an Amazonian people. Columbia University Press.

Balée, W., & Gély, A. (1989). Managed forest succession in Amazonia: The Ka'apor case. Advances in Economic Botany 7, 129-158

Balick, M. J. (1985). Useful plants of Amazonia: A resource of global importance. Chap. 19 in Prance, G.T., and Lovejoy, T.E., eds. Amazonia. Pergamon Press.

Balick, M.J. (1986) Systematics and economic botany of the Oenocarpus-Jessenia (Palmae) complex. Advances in Economic Botany 3: 1-140

Balslev, H., and A. Barfod (1987) Ecuadorean palms- an overview. Opera Botanica 92: 17-35

Balslev, H., and A. Henderson (1987) A New Ammandra ( Palmae) from Ecuador. Systematic Botany 12: 501-504

Balslev, H., C. Grandez, et al. (2008) Useful palms (Arecaceae) near Iquitos, Peruvian Amazon. Rev. peru. biol. 15:121-132

Balslev, H., M. Rios, G. Quezada and B. Nantipa (1997) Palmas útiles en la cordillera de los Huacamayos. PROBONA, Quito

Barriga, H. G. (1974). Flora medicinal de Colombia. Instituto de Ciencias Naturales. Universidad Nacional.

Barriga, R. (1994) Plantas útiles de la Amazonia Peruana: características, usos y posibilidades.. CONCYTEC, Lima

Barrire, P. (1743). Nouuelle Relation de la France Equinoxiale, Paris.

Barros, A. P. et al. (2017) Uso medicinal de plantas na comunidade de Santa Helena, Axixá-Tocantins. Revista Craibeiras de Agroecologia, 1(1).

Beck, H. T. (1990). A survey of the useful species of Paullinia L.(Sapindaceae). Advances in Economic Botany, 41-56.

Beltrán Zapata, G. D. (2016). Conocimiento tradicional y los modos de transmisión de saberes alrededor de las plantas medicinales en la comunidad de Macaquiño (zona Aatiam, territorio del Vaupés) (Master dissertation).

Bennett, B.C., M.A. Baker, and P. Gómez-Andrade (2002) Ethnobotany of the Shuar of Eastern Ecuador. Advances in Economic Botany 14: 1-299

Bentham, G. (1854). On the north Brazilian Euphorbiaceae in the collections of Mr. Spruce. Hooker. Journ. Bot. 6: 368.

Bergman, R. (1980) Economia Amazonica. CAAAP, Lima

Berlin, B., & Berlin, E. A. (1979). Etnobiología subsistencia y nutrición en una sociedad de la selva tropical: los Aguaruna. A. Chirif (comp.), Salud y nutrición en comunidades nativas, 13-47.

Bernal, R. (1992) Colombian palm products. Sustainable harvest and marketing of rain forest products, Washington

Bernal, R., G. Galeano, N. García, I. L. Olivares & C. Cocomá. (2010). Uses and Commercial Prospects for the Wine Palm, Attalea butyracea, in Colombia. Ethnobotany Research and Applications 8: 255-268.

Bertania, S., Bourdy, G., Landaua, I., Robinsonc, J.C., Esterred, P.H. and Deharo, E. (2005). Evaluation of French Guiana traditional antimalarial remedies. Journal of Ethnopharmacology, 98(1-2): 45-54.

Bianchi, C. (1982) Artesanías y técnicas Shuar. Ediciones Mundo Shuar, Quito

Bieski, I. G. C., Leonti, M., Arnason, J. T., Ferrier, J., Rapinski, M., Violante, I. M. P., ... & de Oliveira Martins, D. T. (2015). Ethnobotanical study of medicinal plants by population of valley of Juruena region, legal amazon, Mato Grosso, Brazil. Journal of ethnopharmacology, 173, 383-423.

Bisset, N. G. (1992). Uses, chemistry and pharmacology of Malouetia (Apocynaceae, subf. Apocynoideae). Journal of ethnopharmacology, 36(1), 43-50.

Blohm, H. (1962). Poisonous plants of Venezuela. Poisonous plants of Venezuela. Harvard University Press, Cambridge, Massachusetts.

Bodley, J.H., and F.C. Benson (1979) Cultural ecology of Amazonian palms. Reports of investigations, No. 56. Laboratory of Anthropology, Washington State University, Pullman

Boll, T., J.C. Svenning, J. Vormisto et al. (2005) Spatial distribution and environmental preferences of the piassaba palm Aphandra natalia (Arecaceae) along the Pastaza and Urituyacu rivers in Peru. Forest Ecology and Management 213: 175-183

Boom, B. M. (1990). Useful plants of the Panare Indians of the Venezuelan Guayana. Advances in economic botany, 57-76.

Boom, B.M. (1986) The Chacobo indians and their palms. Principes 30: 63-70

Boom, B.M. (1996). Ethnobotany of the Chácobo Indians, Beni, Bolivia: Second Edition. Advances in Economic Botany , Vol. 4: 1-74

Borchsenius, F., H. Borgtoft, and H. Balslev (1998) Manual to the palms of Ecuador. AAU Reports 37: 1-217

Borgtoft, H. (1992) Uses and management of Aphandra natalia (Palmae) in Ecuador. Bulletin de l´Institute française d´études andines 2: 741-753

Borgtoft, H. (1996) Production and harvest of fibers from Aphandra natalia (Palmae) in Ecuador. Forest Ecology and Management 80: 155-161

Bourdy, G. (1999) Conozcan nuestros árboles, nuestras hierbas. UMSA, CIPTA, IRD. La Paz, Bolivia.

Bourdy, G., C. Valadeau, and J. Albán (2008) Yato` Ramuesh: Plantas Medicinales Yaneshas. PRODAPP-IRD, Lima

Bourdy, G., DeWalt, S. J., De Michel, L. C., Roca, A., Deharo, E., Muñoz, V., ... & Gimenez, A. (2000). Medicinal plants uses of the Tacana, an Amazonian Bolivian ethnic group. Journal of ethnopharmacology, 70(2), 87-109.

Brücher, H. (2012). Useful plants of neotropical origin: and their wild relatives. Springer Science & Business Media.

Brady, G. & H. R. Clauser. (1977). Material handbook. MacGraw-Hill, New York.

Branch, L.C. and Silva, M.F.D. (1983) Folk medicine of Alter do chao, Para, Brazil. Acta Amazonica, 13: 737-797.

Brandão, M.G.L., Grandi, T.S.M., Rocha, E.M.M., Sawyer, D.R. and Krettli, A.U. (1992) Survey of medicinal plants used as antimalarials in the Amazon. Journal of Ethnopharmacology 36(2): 175-182.

Breitbach, U. B., Niehues, M., Lopes, N. P., Faria, J. E., & Brandão, M. G. (2013). Amazonian Brazilian medicinal plants described by CFP von Martius in the 19th century. Journal of ethnopharmacology, 147(1), 180-189.

Brewer-Carias, C., & Steyermark, J. A. (1976). Hallucinogenic snuff drugs of the Yanomamo Caburiwe-Teri in the Cauaburi river, Brazil. Economic Botany, 57-66

Byg, A. & H. Balslev. (2004). Factors affecting local knowledge of palms in Nangaritza valley, Southeastern Ecuador. Journal of Ethnobiology 24: 255-27

Cámara-Leret, R., Fortuna, M. A., & Bascompte, J. (2019). Indigenous knowledge networks in the face of global change. Proceedings of the National Academy of Sciences, 116(20), 9913-9918.

Cárdenas, D. et al. (2007) Plantas útiles y promisorias en la comunidad de Wacurabá (Caño Cuduyarí) en el departamento de Vaupés. Bogotá, SINCHI.

Cárdenas, D., and J.G. Ramírez (2004) Plantas útiles y su incorporación a los sistemas productivos del Departamento del Guaviare ( Amazonia Colombiana). Cadalsia 26: 95-110

Cárdenas, D., and Politis, G.G. (2000) Territorio, movilidad, etnobotánica y manejo del bosque de los Nukak Orientales. Ediciones Uniandes, Santafé de Bogotá

Cárdenas, D., C.A. Marín , L.S. Suárez, et al. (2002) Plantas útiles en dos comunidades del Departamento de Putumayo. Instituto Amazónico de Investigaciones Científicas, SINCHI, Colombia.

Cárdenas, M. (1989) Manual de plantas económicas de Bolivia. Editorial Los Amigos del Libro, La Paz

Caballero-Serrano, V., McLaren, B., Carrasco, J. C., Alday, J. G., Fiallos, L., Amigo, J., & Onaindia, M. (2019). Traditional ecological knowledge and medicinal plant diversity in Ecuadorian Amazon home gardens. Global Ecology and Conservation, 17, e00524.

Cabrera, G., C. Franky, and D. Mahecha (1999) Los nukak: nómadas de la Amazonía colombiana. Unibiblos, Universidad Nacional de Colombia, Bogotá

Cadena-Vargas, C., M. Diazgranados-Cadelo, and H. Bernal-Malagón (2007) Plantas útiles para la elaboración de artesanías de la comunidad indígena Monifue Amena (Amazonas, Colombia). Universitas Scientiarum. Revista de la Facultad de Ciencias 12: 97-116

Califano, M. (1999) Los indios Sirionó de Bolivia oriental. Ciudad Argentina, Buenos Aires

Califano, M., & Distel, A. F. (1982). The Use of a Hallucinogenous Plant among the Mashco (Southwestern Amazonia, Peru). Zeitschrift für Ethnologie, (H. 1), 129-143.

Carod-Artal, F.J. (2012). Curares y timbós, venenos del Amazonas. Revista de neurología 55 (11):689-698.

Castaño-Arboleda, N., D. Cárdenas, and E. Otavo (2007) Ecología, aprovechamiento y manejo sostenible de nueve especies de plantas del departamento del Amazonas, generadoras de productos maderables y no maderables. Instituto amazónico de investigaciones científicas-Sinchi

Castelnau, F. de. (1851) Expedition dans les parlies centrales de l'Amerique du Sud 1843-47. 5: 14, 17, 22-23, 61-62.

Cavalcante, P.B. (1972). Frutas comestíveis da Amazônia I.

Cavalcante, P.B. (1974). Frutas comestíveis da Amazônia II.

Cavalcante, P.B. (1979). Frutas comestíveis da Amazônia III.

CEATA, Trópico, and CI-Bolivia (2007) Transformación del fruto del Majo (Oenocarpus bataua). Recomendaciónes para su aprovechamiento sostenible. Guía para técnicos extensionistas.. CI-Bolivia, La Paz

Cerón, C., and C. Reyes (2007) Aspectos florísticos, ecológicos y etnobotánica de una hectárea de bosque en la comunidad Secoya Sehuaya, Sucumbíos-Ecuador. Caminando sobre el sendero: hacia la conservación del ambiente y la cultura Secoya. Fundación VIHOMA, Quito

Cerón, C., Reyes, C., & Jiménez, E. (2012). Plantas útiles de los Kichwa, centro-norte de la Amazonia ecuatoriana. Cinchonia, 12(1), 22-202.

Cerón, C.E, C. I. Reyes, D. Payaguaje, A. Payaguaje, H. Payaguaje, E. Piaguaje, R. Piaguaje & P. Yépez (2011) Mil y más plantas de la Amazonia ecuatoriana utilizadas por los Secoyas. Cinchonia 11: 13-205.

Cerón, C.E. (1993) Etnobotánica Quichua en la vía Hollín-Loreto, provincia del Napo. Hombre y Ambiente 25: 131-171

Cerón, C.E. (1993) Manejo florístico Shuar-Achuar (Jívaro) del ecosistema amazónico en el Ecuador. Hombre y Ambiente 25: 173-197

Cerón, C.E. (1995) Etnobiología de los Cofanes de Dureno, provincia de Sucumbíos, Ecuador. Museo Ecuatoriano de Ciencias Naturales. Ediciones Abya-Yala, Quito

Cerón, C.E. (2003) Etnobotánica Quichua del Río Yasuní. Cinchonia 4: 1-20

Cerón, C.E., A. Payaguaje, D. Payaguaje et al. (2005) Etnobotánica Secoya. Al inicio del Sendero: Estudios etnobotánicos Secoya. Arboleda, Quito.

Cerón, C.E., and C. Montalvo (2000) Reserva Biológica Limoncocha. Formaciones vegetales, Diversidad y Etnobotánica.. Cinchonia 1: 1-20

Cerón, C.E., and C. Montalvo (2002) Etnobotánica Huaorani de Tivacuno-Tiputini Parque Nacional Yasuni. Cinchonia 3: 64-94

Cerón, C.E., and C.G. Montalvo (1998) Etnobotánica de los Huaorani de Quehueiri-Ono, Napo-Ecuador. Ediciones Abya-Yala, Quito

Cerón, C.E., and C.I. Reyes (2007) Parches de bosque y etnobotánica Shuar en Palora, Morona Santiago-Ecuador.. Cinchonia 8: 66-83

Cerón, C.E., C. Montalvo, C.I. Reyes, and, D. Andi (2005) Etnobotánica Quichua Limoncocha, Sucumbíos-Ecuador. Cinchonia 6: 29-55

Cerón, C.E., C.G. Montalvo, J. Umenda et al. (1994) Etnobotánica y notas sobre la diversidad vegetal en la comunidad Cofán de Sinangüé, Sucumbíos, Ecuador. Ecociencia, Quito

Cerón, C.E., C.I. Reyes, E. Jimenez & J. D. Silva. (2012). Plantas útiles de los Kichwa, centro-norte de la Amazonia ecuatoriana. Cinchonia 12: 22-202.

Cerón, C.E., C.I. Reyes, L. Tonato,A. Grefa et al. (2006) Estructura, composición y etnobotánica del sendero " Ccottacco Shaiqui", Cuyabeno-Ecuador.. Cinchonia 7: 82-114

Cerro, W., W. Nauray, P. Zegarra & E. Flores. (2003). Estudio etnobotánico en las cuencas altas de los ríos Tambopata e Inambari. Proyecto Tambopata Inambari. Pro-Naturaleza, Lima.

Chávez, F. (1996) Estudio preliminar de la familia Arecaceae (Palmae) en el Parque Nacional del Manu (Pakitza y Cocha Cashu). Pp 141-168. In: D. E. Wilson & A. Sandoval (eds). Manu: The biodiversity of southeastern Peru. Editorial Horizonte, Lima.

Chicchon, A. (1992) Chimane resource use and market involvement in The Beni Biosphere Reserve, Bolivia. University of Florida

Cocco, L. (1987). Iyëwei-theri. Quince años entre los Yanomamos. 2nd Edition. Escuela Técnica Don Bosco, Caracas.

Coelho-Ferreira, M. (2009). Medicinal knowledge and plant utilization in an Amazonian coastal community of Marudá, Pará State (Brazil). Journal of Ethnopharmacology, 126(1), 159-175.

Coimbra Jr, C. E. (1985). Estudos de ecologia humana entre os Suruí do Parque Indígena Aripuanã, Rondônia. Plantas de importância econômica. Boletim do Museu Paraense Emílio Goeldi, 2, 37-55.

Coomes, O.T. (2004) Rain forest 'conservation-through-use'? Chambira palm fibre extraction and handicraft production in a a land-constrained community, Peruvian Amazon. Biodiversity and Conservation 13: 351-360

Coomes, O.T. and N. Ban (2004) Cultivated plant species diversity in home gardens of an Amazonian peasant village in northeastern Peru. Economic Botany 58: 420-434

Coomes, O.T., and G.J. Burt (1997) Indigenous market-oriented agroforestry: dissecting local diversity in western Amazonia. Agroforestry Systems 37: 27-44

Copeticona, R.C. (2002) Actividad de recolección de frutos silvestres en las comunidades de San Antonio de Matty y Abaroa (Puerto Rico-Pando). Facultad de Ciencias Puras y Naturales. Universidad Mayor de San Andres, La Paz

Cornejo, M. (1998) Ver, Saber, Poder. Chamanismo de los Yagua de la Amazonía Peruana. IFEA/CAAAP/CAEA-CONICET, Lima

Correa, M. P. (1926). Diccionário das plantas úteis do Brasil. Vols. II (304-309) and VI (229-251). Rio de Janeiro. Inst. Bras. De Desenvolvimento Florestal.

Costa, R. A. (2013). A identidade e o conhecimento etnobotânico dos moradores da Floresta Nacional do Amapá. 2013. 104 f. Dissertação (Mestrado em Biodiversidade Tropical) - Universidade Federal do Amapá, Macapá.

Crevaux, J. N. (1883). Voyages dans l'Amerique du Sud.

Crizón, I. (2001) Por los territorios de la marama. Extracción de la fibra de chiqui chiqui en la amazonía colombiana. Instituto de Estudios Ambientalespara el Desarrollo -IDEADE-, Facultad de Estudios Ambientales y Rurales, Pontificia Universidad Javeriana, Bogotá

Cuatrecasas, J. (1957). The American Species of Dacryodes. Trop. 106: 46-65.

Díaz Piedrahita, S. (1981) Las hojas de las plantas como envoltura de alimentos. Ediciones CIEC, Bogotá

da Conceição Sena, C., Santos, C. D. C. S., da Silva Oliveira, K., & Almeida, R. R. (2019). Análise da comercialização de plantas medicinais no município de Laranjal do Jari-Amapá-Brasil. Revista Eletrônica Casa de Makunaima, 2(4), 105-110

da Costa Moraes, L. L., da Luz Freitas, J., Matos Filho, J. R., Lima, R. B., Borges, C. H. A., & dos Santos, A. C. (2019). A Ethno-knowledge of medicinal plants in a community in the eastern Amazon. Revista de Ciências Agrárias, 42(2), 565-573.

da Costa, I. B., Bonfim, F. P., Pasa, M. C., & Montero, D. A. (2017). Ethnobotanical survey of medicinal flora in the rural community Rio dos Couros, state of Mato Grosso, Brazil. Boletín Latinoamericano y del Caribe de Plantas Medicinales y Aromáticas, 16(1), 53-67.

da Silva Braga, M. D. N., de Morais Batista, D., de Souza, D. B., da Silva Lima, E., de Araújo Pantoja, T. M., Xavier, R. A. T., & Lima, R. A. (2022). Estudo Etnobotânico De Plantas Medicinais Da Família Fabaceae Na Comunidade Cristolândia, Humaitá-Am. Biodiversidade, 21(2).

da Silva, A. F., de Sousa, R. L., Silva, S. G., Costa, J. M., de Albuquerque, L. C. D. S., da Silva Pereira, M. D. G., ... & Cordeiro, Y. E. M. (2021). Etnobotânica de plantas medicinais aromáticas: preparações e usos da flora local em cinco comunidades rurais localizadas na região do Baixo Tocantins, Pará, Brasil. Research, Society and Development, 10(1), e9510111284-e9510111284

Dantas, A. R. et al. (2014). Produção de frutos de urucurizeiros, Attalea excelsa Mart. (Arecaceae), em floresta de várzea no estuário do rio Amazonas. Biota Amazônica, Macapá, v. 4, n. 4, p. 108-114.

Davis, E.W., and J.A. Yost (1983) The ethnobotany of the Waorani of eastern Ecuador. Botanical Museum Leaflets 29: 159-217

Davis, W. (1983) The ethnobotany of Chamairo: Mussatia hyacinthina. Journal of Ethnopharmacology 9: 225-236

De Feo, V. (1991). Uso di piante ad azione antinfiammatoria nell'Alto Ucayali, Perù orientale. Fitoterapia, 62, 481-494.

De Feo, V. (1992). Medicinal and magical plants in the northern Peruvian Andes. FITOTERAPIA-MILANO-, 63, 417-417.

De Feo, V. (1992). Medicinal and magical plants in the northern Peruvian Andes. Fitoterapia-Milano, 63, 417-417.

De Jong, W. (2001) Tree and forest management in the floodplains of the Peruvian Amazon. Forest Ecology and Management 150: 125-134

De la Torre, L., Navarrete, H., Muriel, P., Macía, M. J., & Balslev, H. (2008). Enciclopedia de las Plantas √≤tiles del Ecuador (con extracto de datos). Herbario QCA de la Escuela de Ciencias Biológicas de la Pontificia Universidad Católica del Ecuador & Herbario AAU del Departamento de Ciencias Biológicas de la Universidad de Aarhus.

de Lima Silvaa, W., Aguiarb, G. P. C., Aguiarc, H., Lanad, L. G., Alvese, E. J. A., & Merrittf, B. (2019). Guia etnobotânico de plantas em comunidades Desano (Tukano-oriental) no rio Tiquié-Brasil. In: Cadernos de Etnolingüística, vol. 7, n. 1, p. 1-42

de Menezes Ramos, C. (2018). Segurança alimentar, preservação e conservação ambiental na terra indígena tenharim do marmelos-amazonas, brasil: as plantas e suas utilidades. REDE-Revista Eletrônica do PRODEMA, 11(2), 108-120.

De Oliveira, E. P. B., Peixoto, L. S., Baldissera, M., & Andrighetti, C. R. (2016). Uso, diversidade e conhecimento etnobotânico de plantas medicinais utilizadas para o tratamento da malária no município de nova santa helena-mt. Flovet-.Boletim do Grupo de Pesquisa da Flora, Vegetação e Etnobotânica, 1(8): 89-108

de Oliveira, P. C. (2020). Traditional knowledge of forest medicinal plants of munduruku indigenous people-ipaupixuna. European Journal of Medicinal Plants, 20-35.

de Paula Filho, G. X., Ribeiro, A. F., Moraes, A. F., Penha, W. F., Borges, W. L., & Santos, R. H. S. (2020). Ethnobotanical knowledge on non-conventional food and medicinal plants in Rio Cajari Extractivist Reserve, Amazon, Brazil. https://doi.org/10.21203/rs.3.rs-35316/v3

DeFilipps, Robert A., Shirley L. Maina and Juliette Crepin. (2004). Medicinal Plants of the Guianas (Guyana, Surinam, French Guiana). Smithsonian Institution Press, Washington, D.C.

Denevan, W., and J.M. Treacy (1987) Young managed Fallows at Brillo Nuevo. Advances in Economic Botany 5: 8-49

Descola, P. (1989) La selva culta- Simbolismo y praxis en la ecología de los Achuar. Ediciones Abya-Yala y MLAL, Quito.

Desmarchelier, C., A. Gurni, G. Ciccia & A. M. Giulietti. (1996) Ritual and medicinal plants of the Ese√ïejas of the Amazonian rainforest (Madre de Dios, Peru). Journal of Ethnopharmacology 52: 45-51.

DeWalt, S. J., Bourdy, G., De Michel, L. R. C., & Quenevo, C. (1999). Ethnobotany of the Tacana: quantitative inventories of two permanent plots of northwestern Bolivia. Economic Botany, 53(3), 237-260.

do Nascimento Martinez, L., Pansini, S., Santos, M. G., Silva, D. C., da Silva Rodrigues, F. L., Tada, M. S., & Costa, J. D. A. N. (2018). Avaliação etnobotânica de plantas utilizadas como potenciais antimaláricos na região da Amazônia ocidental brasileira. Interfaces Científicas-Saúde e Ambiente, 6(2), 9-20.

Dos Santos Silva, J. P. G., & Chaves De Oliveira, P. (2016). Etnobotânica de plantas medicinais na comunidade de Várzea Igarapé do Costa, Santarém-Pará, Brasil. Ambiente e Sostenibilidad 6: 136-151

dos Santos, J. X., Reis, A. R. S., Parry, S. M., & de Carvalho, J. C. (2016). Caracterizao Etnobotãnica De Essencias Florestais Com Fins Medicinais Utilizadas Pela Etnia Xipaya, No Municã Pio De Altamira-Pa. Biota Amazônia (Biote Amazonie, Biota Amazonia, Amazonian Biota), 6(2), 1-8.

dos Santos, M. R. A., & de Lima, M. R. (2009). Levantamento dos recursos vegetais utilizados como fitoterápicos no Município de cujubim, Rondônia, Brasil. Boletim de Pesquisa e Desenvolvimento 62.

dos Santos, M. R. A., Lima, M. R. D., & Ferreira, M. D. G. R. (2008). Uso de plantas medicinais pela população de Ariquemes, em Rondônia. Horticultura Brasileira, 26, 244-250.

Dufour, D. L., & Zarucchi, J. L. (1979). Monopteryx angustifolia and Erisma japura: Their Use by Indigenous Peoples in the Northwestern Amazon. Botanical Museum Leaflets, Harvard University, 27(3/4), 69-91.

Dugand, A. (1961) Palms of Colombia. Principes 5: 135-144

Duke, J. A., & Vasquez, R. (1994). Amazonian Ethnobotanical Dictionary. CRC Press, Florida.

Duke, J.A., and Wain, K.K. (1981) Medicinal Plants of the World. Computer index with more than 85,000 entries, 3 vols.

Echeverri, J. A. (2011). Witoto ash salts from the Amazon. Journal of Ethnopharmacology, 138(2), 492-502.

Einzmann, H. (1988) Artesanía indígena del Ecuador: los Cofanes. Revista del CIDAP (Ecuador) 26: 19-86

Elisabetsky, E. and Posey, D.A. (1989) Use of contraceptive and related plants by the Kayapo Indians (Brazil). Journal of Ethnopharmacology, 26(3), pp.299-316.

Ernst, A. (1888). Fischvergiftende Pflanzen. Gesellschaft naturforschender Freunde, Sitzung vom 19. Juni: 111-118

Etter, A.(Ed.) (2001) Puinawai y Nukak. Caracterización Ecológica General de dos Reservas Nacionales Naturales de la Amazonía Colombiana. Instituto de Estudios Ambientales para el Desarrollo (IDEAE), Bogotá

Fanshawe, D. B. (1953) Fish poisons of British Guiana. Kew Bull. 239-240

Farias, J. E. S. (2012). Manejo de açaizais, riqueza florística e uso tradicional de espécies de várzeas do Estuário Amazônico. 102 f. Dissertação (Mestrado em Biodiversidade Tropical) - Universidade Federal do Amapá, Macapá, 2012

Feisal, E. (2009) PFNM en seis comunidades campesinas del norte amazónico boliviano: Causas de éxito o fracaso de comercialización.. PROMAB, Riberalta.

Ferreira De Athayde, S., Mosimann Da Silva, G., Kaiabi, J., Kaiabi, M., Rocha De Souza, H., Ono, K., & Bruna, E. M. (2006). Participatory Research and Management of Arumã (Ischnosiphon Gracilis [Rudge [Köern., Marantaceae) by the Kaiabi People in the Brazilian Amazon. Journal of Ethnobiology, 26(1), 36-59.

Ferreira, A. B., Ming, L. C., Haverroth, M., Daly, D. C., Caballero, J., & Ballesté, A. M. (2015). Plants Used to Treat Malaria in the Regions of Rio Branco-Acre State and Southern Amazonas State-Brazil. International Journal of Phytocosmetics and Natural Ingredients, 2015, 2-9.

Ferrero, P. Andrea. (1966) Los Machiguengas Editorial OPE: Villava-Pamplona. Lima, Peru, p. 193.

Ferreyra, R. (1970) Flora Invasora de los Cultivos de Pucallpi y Tingo Maria.

Finkers, J. (1986). Los Yanomami y su sistema alimentício. Vicariato Apostólico de Puerto Acacucho. Monografía No. 2.

Flores Paitán, S. (1987) Old managed Fallows at Brillo Nuevo. Advances in Economic Botany 5: 53-66

Flores Paitán, S. (1998) Agroforestería amazónica: una alternativa a la agricultura migratoria. Geoecología y desarrollo Amazónico: estudio integrado en la zona de Iquitos, Perú. Annales Universitatis Turkuensis Ser A II 114

Flores, C.F., and P.M.S. Ashton (2000) Harvesting impact and economic value of Geonoma deversa, Arecaceae, an understory palm used for roof thatching in the Peruvian amazon. Economic Botany 54: 267-277

Flores, F. A. (1984). Notes on some medicinal and poisonous plants of Amazonian Peru. Advances in economic botany, 1, 1-8.

Forero, M.C. (2005) Aspectos etnobotánicos de uso y manejo de la familia Arecaceae (palmas) en la comunidad indígena Ticuna de Santa Clara de Tarapoto, del resguardo Ticoya del municipio de Puerto Nariño, Amazonas, Colombia.. Facultad de estudios ambientales y rurales. Pontificia Universidad Javeriana, Bogotá.

Franco, J. (2002). Etnobotanica de la yanchama (Ficus spp: MORACEAE) Amazonas, Colombia. Trabajo de grado. Carrera de Biología. Facultad de Ciencias. Pontificia Universidad Javeriana. Bogota.

Freire, B.H. (2006) Ethnobotany of the Huaorani communities in the Ecuadorian Northwest. Lyonia 10: 7-17

Fuentes, E. (1980). Los Yanomami y las plantas silvestres. Antropológica, 54, 3-138.

Furtado Santos, J.J., Coelho-Ferreira, M. & Costa-Lima, P.G. (2018). Etnobotanica de plantas medicinais em mercados publicos da regiã o metropolitana de belem do parã, brasil. Biota Amazônia (Biote Amazonie, Biota Amazonia, Amazonian Biota), 8(1), 1-9.

Galeano, G. (1992) Las palmas de la región de Araracuara. Tropenbos-Colombia, Bogotá

Galeano, G. & R. Bernal. (2010). Palmas de Colombia. Guía de Campo. Editorial Universidad Nacional de Colombia. Instituto de Ciencias Naturales, Facultad de Ciencias-Universidad Nacional de Colombia, Bogotá.

García Barriga, H. (1974) Flora Medicinal de Colombia. Botánica Médica. Instituto de Ciencias Naturales, Universidad Nacional, Bogotá

García, L.J. , M.J.E. Gómez, S.F.I. Ortiz, and P.J.J. Zuluaga (1996) Principales especies nativas de fauna y flora del Caquetá. Usos actuales y potenciales.. PNR/CORPOICA, Regional Amazonía, Florencia.

Garzón, C., and V. Macuritofe (1992) La noche, las plantas y sus sueños: Aproximación al conocimiento botánico en una cultura amazónica. Corporación Colombiana para la Amazonia, Bogotá

Garzón, N.C. (1985) Aproximación etnobotánica en la comunidad Guayabero de Barrancion-Guaviare. Facultad de Ciencias Humanas. Antropología. Universidad Nacional de Colombia, Bogotá.

Gentry, A. H., & Wettach, R. H. (1986). Fevillea-a new oil seed from Amazonian Peru. Economic Botany, 40(2), 177-185.

Gilmore, M.P., W.H. Eshbaugh and A.M. Greenberg (2002) The use, construction, and importance of canoes among the Maijuna of the Peruvian Amazon. Economic Botany 56: 10-26

Giovannini, P. (2015). Medicinal plants of the Achuar (Jivaro) of Amazonian Ecuador: Ethnobotanical survey and comparison with other Amazonian pharmacopoeias. Journal of ethnopharmacology, 164, 78-88.

Girard, R. (1958) Indios Selváticos de la Amazonia Peruana. Libro Mex, México

Glenboski, L.L. (1983) The Ethnobotany Of The Tukuna Indians. Universidad Nacional de Colombia, Bogota.

Gomez, D., L. Lebrun, N. Paymal, and A. Soldi (1996) Palmas útiles en la provincia de Pastaza. Amazonia ecuatoriana. Manual práctico. Serie Manuales de plantas útiles amazónicas 1. Fundación Omaere. Quito

González-Pérez, S. E., Coelho-Ferreira, M., Robert, P. D., & Garcés, C. L. L. (2012). Conhecimento e usos do babaçu (Attalea speciosa Mart. e Attalea eichleri (Drude) AJ Hend.) entre os Mebêngôkre-Kayapó da Terra Indígena Las Casas, estado do Pará, Brasil. Acta Botanica Brasilica, 26, 295-308

Grández, C.A., and A. Henderson (1993) A new record of Manicaria for Peru. Principes 37: 159-160

Grenand, P. (1980). Introduction d I√ïEtude de l√ïllniuers Wayapi, Tradition Orale No. 40, Societe d√ïEtudes Linguistiques et Anthropologiques de France, Paris.

Guèze, M., Luz, A. C., Paneque-Gálvez, J., Macía, M. J., Orta-Martínez, M., Pino, J., & Reyes-García, V. (2014). Are ecologically important tree species the most useful? A case study from indigenous people in the Bolivian Amazon. Economic Botany, 68, 1-15.

Guallart, J.M. (1968) Nomenclatura Jibaro-Aguaruna de Palmeras en el Distrito de Cenepa.. Biota 7: 230-249

Gumilla, J. (1791). Historia natural, civil y geografica de las naciones situadas en las riveras del rio Orinoco. 2 Vols. (First issued as " El Orinoco Ilustrado . . . ", 1741.)

Gutiérrez-Vásquez, C.A. and R. Peralta (2001) Palmas comunes de Pando, Santa Cruz de la Sierra, Bolivia. PANFOR/OIMT/BOLFOR

Gutiérrez, J. M. B., Ortiz, A. F. L., Perea, E. M., & Méndez, J. J. (2014). Wild medicinal plants used by Colombian Kofan Indians to treat cutaneous leishmaniasis. Revista Cubana de Plantas Medicinales, 19(4), 407-420.

Gutierrez, G. (1943). Estudio sobre los principales barbascos colombianos. Revista Fc. Nac. Agron. Bogotá 7: 77-93

Guyot, M. (1972) La maison des Indiens Bora et Miraña. Journal de la Societe des americanistes 61: 141-176

Guyot, M. (1979) La historia del mar de danta, el Caqueta: una fase de la evolucion cultural en el noroeste amazonico. Journal de la Societe des americanistes 66: 99-123

Hübschmann, L.K., L.P. Kvist, C. Grandez et al. (2007) Uses of Vara Casha - a Neotropical Liana Palm, Desmoncus polyacanthos - in Iquitos, Peru. Palms 51: 167-176

Hajdu, Z., & Hohmann, J. (2012). An ethnopharmacological survey of the traditional medicine utilized in the community of Porvenir, Bajo Paraguá Indian Reservation, Bolivia. Journal of ethnopharmacology, 139(3), 838-857.

Hamlin, C.C., and J. Salick (2003) Yanesha agriculture in the upper Peruvian Amazon: Persistence and change fifteen years down the 'road'. Economic Botany 57: 163-180

Hansson, A., Veliz, G., Naquira, C., Amren, M., Arroyo, M., & Arevalo, G. (1986). Preclinical and clinical studies with latex from Ficus glabrata HBK, a traditional intestinal anthelminthic in the Amazonian area. Journal of Ethnopharmacology, 17(2), 105-138.

Harner, M. J. (1984) The Jívaro. People of the Sacred Waterfalls. University of California Press, Berkeley

Hartwell, J. L. (1982). Plants used against cancer: a survey. Quarterman Publications, Inc. Lawrence, MA. 710 pp.

Haverroth, M., Negreiros, P. R. M., & Barros, L. C. P. (2010). Ethnobiology and health among the Kulina people from the Upper Envira River, State of Acre, Brazil. The Open Complementary Medicine Journal, 2(1)

Hegnauer, R. (1964). Chemotaxonomie der Pflanzen-Birkhauser Verlag. Basel. Vols. 1-6

Heizer, R. F. (1953). Aboriginal fish poisons. US. American Ethnology Bureau Bulletin. 151 No 38: 231-283

Henderson, A., and F. Chávez (1993) Desmoncus as a useful palm in the western Amazon basin. Principes 37: 184-186

Henkemans, A. (2001) Tranquilidad and Hardship in the Forest: Livelihoods and Perceptions of Camba Forest dwellers in the northern Bolivian Amazon. PROMAB, Riberalta

Hibert, F., Sabatier, D., Andrivot, J., Scotti-Saintagne, C., Gonzalez, S., Prévost, M. F., ... & Richard-Hansen, C. (2011). Botany, genetics and ethnobotany: a crossed investigation on the elusive tapir's diet in French Guiana. PLoS One, 6(10), e25850.

Hinojosa, I. (1991) Plantas utilizadas por los Mosetenes de Santa Ana (Alto Beni, Depto. La Paz).. Facultad de Ciencias Puras y Naturales. Universidad Mayor de San Andres, La Paz.

Hinojosa, I., E. Uzquiano, E., and J. Flores (2001) Los Yuracaré: su conocimiento, experiencia y la utilización de recursos vegetales en el río Chapare. Instituto de Ecología, Universidad Mayor de San Andrés, La Paz

Hissink, K., and A. Hahn (2000) Los Tacana- datos sobre la historia de su civilización. Plural Editores, La Paz

Hoehne, F.C. (1937). Botanica e Agricultura no Brasil no seculo XVI. Bibl. Pedag. Brasil., V. Brasiliana. 71

Hoffman, B. & Ruysschaert, S. (2017). Lianas of the Guianas: a field guide to woody climbers in the tropical forests of Guyana, Suriname and French Guiana. LM Publishers.

Holm-Jensen, O. and H. Balslev (1995) Ethnobotany of the Fiber Palm Astrocaryum Chambira ( Arecaceae) in Amazonian Ecuador. Economic Botany 49: 309-319

Holmberg, A.R. (1978) Nómadas del arco largo: los Sirionó del oriente boliviano. Instituto Indigenista Interamericano, México

Horackova, J., Chuspe Zans, M. E., Kokoska, L., Sulaiman, N., Clavo Peralta, Z. M., Bortl, L., & Polesny, Z. (2023). Ethnobotanical inventory of medicinal plants used by Cashinahua (Huni Kuin) herbalists in Purus Province, Peruvian Amazon. Journal of Ethnobiology and Ethnomedicine, 19(1), 16.

Howes, F. N. (1930). Fish-poison plants. Kew Bulletin 4, 129-153

Huanca, T. (1999) Tsimane Indigenous Knowledge, Swidden Fallow Management and Conservation.. University of Florida, Gainesville

Huertas, B. (2007) Nuestro territorio Kampu Piyawi (Shawi). Terra Nuova, Lima

Humboldt, A. von & A. Bonpland. (1807). Voyage aux régions equinoxiales du Nouveau Continent (Relation historique du voyage) 2 : 547-556

Iglesias, G. (1987) Hierbas medicinales de los Quichua del Napo. Abya-Yala, Quito

Iglesias, G. (1989) Hierbas medicinales de los Quichua del Napo. Enfermedades femeninas y enfermedades del "susto". Abya-Yala, Quito

Iglesias, G. (1989) Sacha Jambi- El uso de las plantas en la medicina tradicional de los Quichuas del Napo. Ediciones Abya-Yala, Quito

Irvine, D. (1989) Succesion Management and Resource Distribution in an Amazonian Rain Forest. Advances in Economic Botany 7: 223-237

Játiva, V. & R. Alarcón. (1994). Sobre la etnobotánica y la comercialización de la ungurahua, Oenocarpus bataua (Arecaceae), en la zona del Alto Napo, Ecuador. Pp 53-89. In: R. Alarcón, P. A. Mena, & A. Soldi (eds). Etnobotánica, valoración económica y comercialización de recursos florísticos silvestres en el Alto Napo. Ecuador. Ecociencia, Quito.

Jernigan, K. A. (2009). Barking up the same tree: a comparison of ethnomedicine and canine ethnoveterinary medicine among the Aguaruna. Journal of Ethnobiology and Ethnomedicine, 5(1), 1-9.

Jobert, C. (1878). Sur la preparation du Curare. Compt. Rend. Acad. Sci. Paris 86: 121-122.

Johnson, D. (1975) Some palm products of the Peruvian Amazon. Principes 19: 78-79

Johnson, D., and K. Mejía (1998) The Making of a Dugout Canoe from the Trunk of the Palm Iriartea deltoidea. Principes 42: 201-205,208

Jordan, C.B. (1970) A study of germination and use in twelve palms of Northeastern Peru. Principes 14: 26-32

Kahn, F.,and K. Mejía (1987) Notes on the biology, ecology, and use of a small Amazonian palm: Lepidocaryum tessmannii. Principes 31: 14-19

Kainer, K.A. and Duryea, M.L. (1992). Tapping women√ïs knowledge: plant resource use in extractive reserves, Acre, Brazil. Economic Botany 46(4): 408-425.

Kamen-Kaye, D. (1977). Ichthyotoxic Plants and the Term" barbasco". Botanical Museum Leaflets, Harvard University, 25(2), 71-90.

Karsten, R. (1988) La vida y la cultura de los Shuar. Tomo I. Abya-Yala, Quito

Kerharo, H., Guichard, F. & Bouquet, A. (1959-1961). Les végétaux ichtyotoxiques (poison de peche). Bull, et Mém. Fac. Med. Pharm. Dakar. 8: 314-329: 9: 355-386: 10: 223-242.

Kermath, B. M., Bennett, B. C., & Pulsipher, L. M. (2018). Food Plants in the Americas.

Kerr, W. E., Posey, D. A., & Wolter Filho, W. (1978). Cupá, ou cipó-babão, alimento de alguns índios amazônicos. Acta Amazonica, 8, 702-705.

Kffuri, C. W. (2014). Etnobotânica de plantas antimaláricas em comunidades indígenas da região do Alto Rio Negro-Amazonas-Brasil. PhD Thesis.

Killip, E.P. & Smith, A.C. (1935) Some American plants used as fish poisons. Bureau of Plant Industry. USDA.

Kronik, J. et al. (1999) Fééjahisuu. Palmas de los Nietos de la Tierra y Montaña Verde del Centro. Centro de Investigación y Desarrollo, Copenhague

Krukoff, B. A. (1965). Supplementary notes on the American species of Strychnos VII. N.Y. Bot. Gard. 12(2): 7, 42-43.

Krukoff, B. A., & Moldenke, H. N. (1938). Studies of American Menispermaceae, with special reference to species used in preparation of arrow-poisons. Brittonia, 3(1), 1-74.

Krukoff, B. A., & Monachino, J. (1942). The American species of Strychnos. Brittonia 4: 248-332

Krukoff, B. A., & Monachino, J. (1946). The genus Strychnos in Venezuela. Darwiniana, 7(2), 185-193.

Krukoff, B. A., & Monachino, J. (1947). Supplementary Notes on the American Species of Strychnos IV. Bol. Tec. Inst. Agron. Belem 11: 3-15

Krukoff, B. A., & Smith, A. C. (1937). Notes on the botanical components of curare. Bulletin of the Torrey Botanical Club, 401-409.

Krukoff, B. A., & Smith, A. C. (1937). Rotenone-yielding plants of South America. American Journal of Botany, 573-587.

Krukoff, B. A., & Smith, A. C. (1939). Notes on the botanical components of curare-II. Bulletin of the Torrey Botanical Club 66, 305-314.

Kvist, L. P., Oré-Balbín, I. C., & Llapapasca-Samaniego, D. C. (1998). Plantas utilizadas en trastornos ginecológicos, parto y control de natalidad en mujeres de la parte baja del río Ucayali, Amazonas Peruana. Folia Amazónica, 9(1-2), 115-141.

Kvist, L., M.K. Andersen, J. Stagegaard, M. Hesselsoe, and C. Llapapasca (2001) Extraction from woody plants in flood plain communities in Amazonian Peru: use, choice, evaluation and conservation status of resoruces. Forest Ecology and Management 150: 147-174

Lévi-Strauss, C. (1952). The use of wild plants in tropical South America. Economic Botany, 6(3), 252-270.

López Camacho, R. (2006) Manual de identificación de especies no maderables del corregimiento de Tarapacá, Colombia. Bogota, Sinchi

López-Parodi, J. (1988) The use of palms and other native plants in non-conventional, low cost rural housing in the Peruvian Amazon. Advances in Economic Botany 6: 119-129

López, R., D. Cárdenas & C. Marín. (1998). Plantas de Uso Potencial (no maderable) en el Norte del Departamento del Guaviare, Amazonía Colombiana. The Field Museum and The Andrew Mellon Foundation, Chicago.

La Rotta, C. (1983) Observaciones etnobotánicas sobre algunas especies utilizadas por la comunidad indígena Andoque (Amazonas, Colombia). Universidad de Colombia, Bogotá.

La Rotta, C., P. Miraña, M. Miraña, B. Miraña, M. Miraña & N. Yucuna. (1987). Estudio etnobotánico sobre las especies utilizadas por la comunidad indígena Miraña, Amazonas, Colombia. World Wildlife Fundation, Bogotá.

La Rotta, C., P. Miraña, M. Miraña, B. Miraña, M. Miraña & N. Yucuna. (1989). Especies utilizadas por la Comunidad Miraña. Estudio Etnobotánico. Fondo para la protección del medio ambiente √íJosé Celestino Mutis√ì, FEN Colombia. Bogotá.

Lambda, SS. (1970) Indian piscidal plants. Econ. Bot. 24: 134-136

Langevin, M. (2002) Mapajo. Nuestra selva y cultura. PRAIA (FIDA/CAF), La Paz.

Le Cointe, P. (1947) √ßrvores e plantas úteis:(indígenas e aclimadas). Brasiliana, vol. 251. Rio ; 506 p. il.

Lewis, W. H., & Elvin-Lewis, M. P. (1977). Medical botany: plants affecting human health. John Wiley & Sons, NY. 515 pp.

Lima, R. B., Souto, R. N. P., & de Medeiros, F. A. (2019). Espécies vegetais usadas como repelentes e inseticidas no estado do Amapá, BR. Revista Brasileira de Agroecologia, 14(3), 14-14.

Lizot, J. (1984). Les Yanõmami centraux. Cahiers de l√îHomme, Éditions de L√îEHESS, Paris

Lozano Balcázar, A. (2005). Los barbascos utilizados por los Ticuna del PNN Amacayacu (Bachelor's thesis, Uniandes).

Luna, L. E. (1982). El concepto de plantas que enseñan, entre los cuatro Shamanes mestizos de Iquitos nordeste del Perú. Revista Colombiana de Antropología, 24, 45-66.

Luziatelli, G., M. Sørensen, I. Theilade & P. Mølgaard. (2010). Asháninka medicinal plants: a case study from the native community of Bajo Quimiriki, Junín, Peru. Journal of Ethnobiology and Ethnomedicine 6: 21.

Macía, M. J. (2004). A comparison of useful pteridophytes between two Amerindian groups from Amazonian Bolivia and Ecuador. American Fern Journal, 94(1), 39-46.

Macía, M.J. (2004) Multiplicity in palm uses by the Huaorani of Amazonian Ecuador. Botanical Journal of the Linnean Society 144: 149-159

Macía, unpubl.

Macbride, J.F. (1960) Flora of Peru Vol. XIII Part 1 Nº 2. Field Museum of Natural History, Chicago.

Macbridge, F. (1956). Flora of Peru. Field Mus. Nat. Hist. Publ., Bot. Ser., Vol. 13 (3a) No. 2: 291-744

MacBridge, J.F. (1936-). Flora of Perú. Field Museum of Natural History, Botanical Services, Chicago.

Magnin, J., Breve Description de la Provincia de Quito (1740). In C. Bayle, Descubridores Jesuitas de1 Amazonas, Instituto Gonzalo Fernandez de Oviedo, Madrid, 1940, p. 46.

Maia, A. C., Queiroz, T. D. S., de Oliveira, A. E., Matos, M. V., & Arcanjo-Silva, S. (2020). Fitoterapia Familiar no Assentamento Madre Cristina (Ariquemes, Rondônia). Brazilian Journal of Development, 6(11), 89780-89798

Marles, R.J., D.A. Neill, and N.R. Farnsworth (1988) A contribution to the ethnopharmacology of the lowland Quichua people of Amazonian Ecuador. Revista de la Academia Colombiana de Ciencias Exactas, Físicas y Naturales 16: 111-120

Marques, W. P. G., dos Anjos, T. O., & da Costa, M. N. R. F. (2020). Plantas medicinais usadas por comunidades ribeirinhas do Estuário Amazônico. Brazilian Journal of Development, 6(10), 74242-74261

Martin, Franklin W., Carl W. Campbell, and Ruth M. Ruberte_. 1987. Perennial Edible Fruits of the Tropics. Agriculture Handbook #642. USDA, Washington, D.C.

Martins, J. E. C. (1989). Plantas medicinais de uso na Amazônia. 2a ediçao.

Martius, C. F. yon. (1830). Ueber die Bereitung des Pfeilg'ftes Urari bet den Indianern Juris am Rio Japura in Nordbrasilien. Buehner, Repert. Pharmacie 36: 340-349.

Martius, C.F. P. von. (1843). Systema Materiae Medicae Vegetabilis Brasiliensis. F. Fleischer. Leipzig. P.67

Martyn, E.B. & Follett Smith, R.R. (1936) The fish poison plants of British Guiana, with special reference to the genera Tephrosia and Lonchocarpus. Agric. J. British Guiana 7: 154-159

Mata, N. D. S. Participação da mulher waiãpi no uso tradicional de plantas medicinais. (2009). 141 f. Dissertação (Mestrado em Desenvolvimento Regional) - Universidade Federal do Amapá, Macapá

Maxwell, N. (1990). Witch doctor's apprentice: hunting for medicinal plants in the Amazon. 3rd edition. Citadel Press, New York. 391 pp.

Mayer, W. (2006) THE PIASSABA PALM: CONSERVATION AND DEVELOPMENT IN THE BUFFER ZONE OF PERU√ïS CORDILLERA AZUL NATIONAL PARK. Nicholas School of the Environment and Earth Sciences, Duke University

Medicinal Plants of the Guianas

Mejía, K. (1988) Utilization of palms in eleven Mestizo villages of the Peruvian Amazon (Ucayali river, department of Loreto). Advances in Economic Botany 6: 130-136

Mejía, K. (1992) Las palmeras en los mercados de Iquitos. Bulletin de l´Institute française d´études andines 21: 755-769

Mejía, K., and E. Rengifo (2000) Plantas Medicinales de Uso Popular en la Amazonía Peruana. Agencia Española De Cooperación Internacional (Aeci) Y El Instituto De Investigaciones De La Amazonía Peruana (IIAP), Lima

Mejía, K.M. (1983) Palmeras y el selvícola amazónico. Museo Historia Natural, Universidad Nacional Mayor de San Marcos, Lima.

Mejía, L. E., & Turbay, S. (2009). Los venenos de cacería en la Amazonia colombiana:√Ä sustancias letales o fuente de vitalidad?. Boletín de Antropología Universidad de Antioquia, 23(40), 129-153.

Melo, N. C. (2015). Avaliação da atividade protetora solar in vitro das espécies pau-mulato (Calycophyllum spruceanum (Benth.) Hook. f. ex K. Schum) e ipê-amarelo (Tabebuia aurea (Silva Manso) Benth. & Hook. f. ex S. Moore). 2015. 90 f. Dissertação (Mestrado em Ciências da Saúde) - Universidade Federal do Amapá, Macapá

Mendoza, D.E. and A. Panduro (2005) El tejido de las hojas de palmera en la vivienda amazónica. Proyecto Araucaria Amazonas Nauta/ Agencia Española de Cooperación internacional - Gobierno Regional de Loreto - Iquitos, Perú

Mendoza, N. U. (1961). Algunos colorantes vegetales usados por las tribus indígenas de Colombia. Revista Colombiana de Antropología, 10, 333-340.

Mendoza, P. (1994) Identificación de los frutos comestibles silvestres recolectados por los indígenas Huaorani de la comunidad de Toñiampari en la Amazonía del Ecuador. Pontificia Universidad Católica del Ecuador, Quito

Meneguelli, A.Z. (2020) Ethnopharmacological and botanical evaluation of medicinal plants used by Brazilian Amazon Indian community. Interacoes 31(3), 633-645

Mesa, C. L. (2011). Etnobotánica de Palmas en la Amazonia Colombiana: Comunidades Indígenas Piapocos del río Guaviare, como estudio de caso. Master thesis. Línea Manejo y Conservación de Vida Silvestre. Universidad Nacional de Colombia, Facultad de Ciencias, Departamento de Biología. Bogotá.

Mesa, L., & Galeano, G. (2013). Palms uses in the Colombian Amazon. Caldasia, 35(2), 351-369.

Miller, C. (2002) Fruit production of the ungurahua palm (Oenocarpus bataua subsp bataua, Arecaceae) in an indigenous managed reserve. Economic Botany 56: 165-176

Miller, R. P., Wandelli, E. V., & Grenand, P. (1989). Conhecimento e utilização da floresta pelos índios Waimiri-Atroari do Rio Camanau-Amazonas. Acta Botanica Brasilica, 3, 47-56

Milliken, W. (2021). Traditional Medicines Amongst Indigenous Groups in Roraima, Brazil: A Retrospective. Ethnoscientia-Brazilian Journal of Ethnobiology and Ethnoecology, 6(3), 116-139.

Milliken, W., Albert, B., Gomez, G.G. (1999). Yanomami: a forest people. Vol. 701. Royal Botanic Gardens, Kew.

Milliken, W., Walker, B. E., Howes, M. J. R., Forest, F., & Lughadha, E. N. (2021). Plants used traditionally as antimalarials in Latin America: mining the Tree of Life for potential new medicines. Journal of Ethnopharmacology, 114221.

Mitchell, John D. (1992). Additions to Anacardium (Anacardiaceae). Anacardium amapaënse, a New Species from French Guiana and Eastern Amazonian Brazil. Brittonia 44(3):331-338.

Mollinedo, L. G. (2000). Motacú (Attalea phalerata) en la Comunidad Leco Irimo. Publicaciones Proyecto de Investigación CIDOB-DFID, Santa Cruz.

Monachino, J. (1949). A Revision of Ryania (Flacourtiaceae). Lloydia 12(1): 1-29

Mondragón, M.L., and R. Smith (1997) Bete Quiwiguimamo. Salvando el bosque para vivir sano. Ediciones Abya-Yala, Quito

Moore, S.J., N. Hill, C. Ruiz, and M.M. Cameron (2007) Field evaluation of traditionally used plant-based insect repellents and fumigants against the malaria vector Anopheles darlingi in Riberalta, Bolivian Amazon. Journal of Medical Entomology 44: 624-630

Moraes, M. (2004) Flora de palmeras de Bolivia. Plural Editores, La Paz

Moraes, M., and J. Sarmiento (1999) La jatata (Geonoma deversa (Poit.) Kunth, Palmae)- un ejemplo de producto forestal forestal no maderable en Bolivia: uso tradicional en el este del departamento de La Paz.. Revista de la Sociedad Boliviana de Botánica 2: 183-196

Moraes, M., F. Borchsenius, et al. (1996) Notes on the biology and uses of the Motacú palm (Attalea phalerata, Arecaceae) from Bolivia. Economic Botany 50: 423-428

Moraes, M., J. Sarmiento,and E. Oviedo (1995) Richness and uses in a diverse palm site in Bolivia. Biodiversity and Conservation 4: 719-727

Morcote-Ríos, G., G. Cabrera-Becerra et al. (1998) Las palmas entre los grupos cazadores-recolectores de la Amazonia colombiana. Cadalsia 20: 57-74

Moreno Suárez, L., and O.I. Moreno Suárez (2006) Colecciones de las palmeras de Bolivia. Editorial FAN, Santa Cruz de la Sierra

Moretti, C., & Grenand, P. (1982). Les nivrées ou plantes ichtyotoxiques de la Guyane française. Journal of Ethnopharmacology, 6(2), 139-160.

Mori, S.A. (1979) In: G.T. Prance & S.A. Mori, Lecythidaceae. I. Fl. Neotrop. Monogr. 21: 128-197

Mors, W.B., Nascimento, M.C., Valle, J.R. & Aragao, J.A. (1973) Ichthyotoxic activity of plants of the genus Derris and compounds isolated therefrom. Cienc. e Cult. 25: 647-648

Muñoz, V., Sauvain, M., Bourdy, G., Callapa, J., Bergeron, S., Rojas, I., ... & Deharo, E. (2000). A search for natural bioactive compounds in Bolivia through a multidisciplinary approach: Part I. Evaluation of the antimalarial activity of plants used by the Chacobo Indians. Journal of Ethnopharmacology, 69(2), 127-137.

Mundo Shuar (1977) Las plantas. Mundo Shuar 5

Nascimento, E. S. (2011). Levantamento dos conhecimentos etnobotânicos de comunidades ribeirinhos do estuário amapaense. 86 f. Trabalho de Conclusão de Curso (Graduação em Engenharia Florestal) - Universidade do Estado do Amapá, Macapá.

Nimuendaju, C. (1956) Os Apinayé. Bol. Do Mus. Emilio Goeldi 12: 1-150. Pg. 69.

Odonne, G., Berger, F., Stien, D., Grenand, P., & Bourdy, G. (2011). Treatment of leishmaniasis in the Oyapock basin (French Guiana): A KAP survey and analysis of the evolution of phytotherapy knowledge amongst Wayãpi Indians. Journal of Ethnopharmacology, 137(3), 1228-1239.

Odonne, G., Valadeau, C., Alban-Castillo, J., Stien, D., Sauvain, M., & Bourdy, G. (2013). Medical ethnobotany of the Chayahuita of the Paranapura basin (Peruvian Amazon). Journal of Ethnopharmacology, 146(1), 127-153.

Ojeda, P. (1994) Diagnóstico etnobotánico y comercialización del morete, Mauritia flexuosa (Arecaceae), en la zona del Alto Napo, Ecuador. Etnobotánica, valoración económica y comercialización de recursos florísticos silvestres en el Alto Napo, Ecuador. Ecociencia. Quito.

Oliveira, S. K. S. (2016). Etnobotânica em duas comunidades da terra indígena São Marcos. Roraima, Brasil (Doctoral dissertation, Tese (Doutorado em Biodiversidade e Conservação)-Instituto de Ciências Biológicas, Universidade Federal do Pará. Belém, 2016, 113f).

Oré, I., and D. Llapapasca (1996) Huertas Domesticas Como Sistema Tradicional De Cultivo En Moena Caño, Rio Amazonas, Iquitos - Peru. Folia Amazonica 8: 91-110

Ordinaire, 0. (1892) Du Pacifique a l√ïAtlantique par les Andes Peruviennes et l√ïAmazone. Plon, Nourrit & Co., Paris, pp. 131-133.

Ortiz, R. (1994) Uso, conocimiento y manejo de lagunos recursos naturales en el mundo Yucuna (Mirití-Paraná, Amazonas, Colombia).. Ediciones Abya-Yala, Quito

Otterburg, M., and M. Mamani (2008) Buenas prácticas de aprovechamiento de Jatata. Tropico, La Paz

Pérez Arbeláez, E. (1956). Plantas útiles de Colombia: ensayo de botánica colombiana aplicada. Librería Colombiana, Bogotá.

Pérez, D. (2002) Etnobotánica medicinal y biocidas para malaria en la Región de Ucayali. Folia Amazonica 13: 87-108

Pérez, H., P.E. Bucheli, and B.G. Benavides (2006) La agroforestería en Guainía: Una alternativa sostenible. Instituto amazónico de investigaciones científicas-Sinchi, Bogotá

Pacheco, T., R. Burga, P.A. Angulo, and J. Torres (1998) Evaluación de Bosques Secundarios de la zona de Iquitos. Geoecología y desarrollo Amazónico: estudio integrado en la zona de Iquitos, Perú. Annales Universitatis Turkuensis Ser A II 114

Padoch, C. (1988) Aguaje (Mauritia flexuosa L.f.) in the economy of Iquitos, Peru. Advances in Economic Botany 6: 214-224

Padoch, C., and W. De Jong (1989) Production and profit in agroforestry: an example from the Peruvian Amazon. Fragile lands of Latin America. Strategies for sustainable development. Westview Press, London

Padoch, C., J. Chota Inuma, W. De Jong, and J. Unruh (1987) Market-Oriented Agroforestry at Tamshiyacu. Advances in Economic Botany 5: 90-96

Padoch, C., J.Chota-Inuma, W. De Jong,and J. Unruh (1985) Amazonian agroforestry: a market-oriented system in Peru. Agroforestry Systems 3: 47-58

Paniagua Zambrana et al., unpubl.

Paniagua Zambrana, N.Y. (1998) Estudio comparativo de la densidad y los niveles de producción de hojas, frutos y semilas en poblaciones naturales del Attalea phalerata ( Palmae) sometidas a diferente intensidad de extracción (Riberalta, Depto. Beni, Bolivia). Universidad Mayor de San Andrés

Paniagua Zambrana, N.Y. (2001) Guía de plantas útiles de la comunidad de San José de Uchupiamonas. FUND-ECO/LIDEMA/Herbario Nacional de Bolivia. La Paz

Paniagua Zambrana, N.Y. (2005) Diversidad, densidad, distribución y uso de las palmas en la región del Madidi, noreste del departamento de La Paz (Bolivia). Ecología en Bolivia 40: 265-280

Paniagua-Zambrana, N. (2005) Conocimiento y uso de las palmas del Abanico del ri_o Pastaza en la Amazonía Peruana. Projekt Arbejde. Dept. of Systematic Botany, Institute of Biology, University of Aarhus, Aarhus

Paniagua-Zambrana, N., Cámara-Leret, R., & Macía, M. J. (2015). Patterns of medicinal use of palms across northwestern South America. The Botanical Review, 81, 317-415.

Parrotta, J. A., Francis, J. K., & de Almeida, R. R. (1995). Trees of the Tapajós: a photographic field guide. Trees of the Tapajós: A photographic field guide.

Peña-Claros, M. (1996) Ecology and socioeconomics of palm heart extraction from wild populations of Euterpe precatoria Mar. in eastern Bolivia.. University of Florida, Gainesville

Peñuela Salazar, M.L. (2002). Estudio etnobotánico del genero Brosimum Sw. Moraceae y su potencial de uso en Leticia y Araracuara, amazonia colombiana. Tesis, UNAL

Peckolt, T. (1910) Heil- und Nutzpflanzen Brasiliens. Berichfe der Deuischen Pharmazeutischen Geseiischaft 20, 36-58, 43.

Pedrollo, C. T., Kinupp, V. F., Shepard Jr, G., & Heinrich, M. (2016). Medicinal plants at Rio Jauaperi, Brazilian Amazon: Ethnobotanical survey and environmental conservation. Journal of ethnopharmacology, 186, 111-124.

Pennington, T. D. (1990). Flora neotropica. Monograph 52. Sapotaceae. New York Botanical Garden for the Organization for Flora Neotropica.

Pereira, L. A. et al. (2007). Plantas medicinais de uma comunidade quilombola na Amazônia Oriental: aspectos utilitários de espécies das famílias Piperaceae e Solanaceae. Rev. Bras. de Agroecologia, v. 2, n. 2, p. 1385-1388.

Peters, C.M., A.H. Gentry, ,and R. Mendelsohn (1989) Valuation of an Amazonian rainforest. Nature 339: 655-656

Pezzuti, J., & Chaves, R. P. (2009). Etnografia e manejo de recursos naturais pelos índios Deni, Amazonas, Brasil. Acta Amazonica, 39, 121-138.

Phillips, O.L. (1993) The potential for harvesting fruits in tropical rainforests: new data from Amazonian Peru. Biodiversity and Conservation 2: 18-38

Pilnik, M. S., Argentim, T., Kinupp, V. F., Haverroth, M., & Ming, L. C. (2023). Traditional botanical knowledge: food plants from the Huni Ku_ indigenous people, Acre, western Brazilian Amazon. Rodriguésia, 74, e00482021.

Pinedo-Vasquez, M., D. Zarin, P. Jipp, et al. (1990) Use-Values of Tree Species in a Communal Forest Reserve in Northeast Peru. Conservation Biology 4: 405-416

Pinkley, H. V. (1969). Plant admixtures to ayahuasca, the South American hallucinogenic drink. Lloydia, 32(3), 305-314.

Pinkley, H. V. (1973). The ethno-ecology of the Kofan Indians (Doctoral dissertation, Harvard University).

Pinto, A. A. D. C., & Maduro, C. B. (2003). Produtos e subprodutos da medicina popular comercializados na cidade de Boa Vista, Roraima. Acta Amazônica, 33, 281-290.

Pipoly, J.J. (1983) Contributions toward a monograph of Cybianthus (Myrsinaceae): III. A revision of subgenus Laxiflorus. Brittonia 35: 61-80

Pires, J. M. (1978) Plantas ichthyotoxicas: Aspecto da botânica systematica. Ciencia e Cult. Supl.

Pittier, H. (1926). Manual de las plantas usuales de Venezuela. Litografia del comercio.

Plotkin, M. J. (1994). Tales of a Shaman's Apprentice: An Ethnobotanist Searches for New Medicines in the Rain Forest. Viking Press, NY. 318 pp.

Plowman, T. (1977). Brunfelsia in ethnomedicine.√äBotanical Museum Leaflets, Harvard University,√ä25(10), 289-320.

Plowman, T. (1980) Chamairo: Mussatia hyacinthina - an admixture to coca from Amazonian Peru and Bolivia. Botanical Museum Leaflets, Harvard Univesrity 28: 253-261.

Plowman, T. (1981) Amazonian coca, J Ethnopharmacology 3: 195-225

Plowman, T. C., Leuchtmann, A., Blaney, C., & Clay, K. (1990). Significance of the fungus Balansia cyperi infecting medicinal species of Cyperus (Cyperaceae) from Amazonia. Economic Botany, 452-462.

Polesna, L., Polesny, Z., Clavo, M. Z., Hansson, A., & Kokoska, L. (2011). Ethnopharmacological inventory of plants used in coronel Portillo province of Ucayali department, Peru. Pharmaceutical Biology, 49(2), 125-136.

Ponce, M. (1992) Etnobotánica de palmas de Jatun Sacha. Pages 43-51.Memorias del 3er Simposio Colombiano de Etnobotánica. INCIVA.

Posey, D. A. (1984). A preliminary report on diversified management of tropical forest by the Kayapo Indians of the Brazilian Amazon. Advances in Economic Botany, 1, 112-126.

Posey, D. A. (2002). Kayapó Ethnoecology and Culture (Vol. 6). Routledge.

Poveda, L.J. (1985-86). Marvels of our Medicinal Flora (in Spanish). Biocenosis Vols. 1 and 2.

Prada Pedreros, S. (1987). Acercamiento etnopiscicolas con los indios ticuna del Parque Nacional Natural Amacayacu, Amazonas (Colombia). Tesis de pregrado, Facultad de Ciencias, UNAL, Bogota. Pg. 56

Prado, M. L. (2008). Las palmas en la comunidad Tikuna de San Martín de Amacayacu: Conocimiento y Uso. Graduate thesis. Maestría en Estudios Amazónicos. Universidad Nacional de Colombia. Sede Amazonia. Leticia, Colombia.

Prance, G. T. (1972). An ethnobotanical comparison of four tribes of Amazonian Indians. Acta amazónica, 2(2), 7-27.

Prance, G. T. (1972). Ethnobotanical notes from Amazonian Brazil. Economic Botany, 26(3), 221-237.

Prance, G. T., Campbell, D. G., & Nelson, B. W. (1977). The ethnobotany of the Paumarí Indians. Economic Botany, 31(2), 129-139.

Proctor, P., J. Pelham, B. Baum, C. Ely, M.A. Rogríguez-Girones (1992) Expedición de la Universidad de Oxford a Bolivia. Investigación etnobotánica de las Palmae en el noroeste del departamento de Pando. 25 junio-7 de septiembre de 1992. Informe final: Sección B.. Universidad de Oxford

Quintana Arias, R. F. (2012). Estudio de plantas medicinales usadas en la comunidad indígena Tikuna del alto Amazonas, Macedonia. Nova, 10(18), 181-193.

Quintana, G., and L. Vargas (1995) Guia popular de plantas utilizadas por los Mosetenes de Covendo, Santa Ana y Muchanes (Alto Beni, Bolivia). FONAMA, La Paz

Ramos, R. S. (2014). Estudo fitoquímico da atividade microbiológica de citoxicidade e larvicida dos óleos essenciais de espécies da família Lamiacea (LAMIALES). 2014. 86 f. Dissertação (Mestrado em Ciências Farmacêuticas) - Universidade Federal do Amapá, Macapá.

Ramos, R. S. et al. (2015). Estudo físico-químico e avaliação do potencial larvicida do extrato etanólico das cascas do caule de Licania macrophylla Benth. Biota Amazônica, Macapá, v. 5, n. 1, p. 74-78.

Remy, F.E. (1908) Apuntes sobre el Clima y Flora de la Region de1 Pichis. In: Carlos Larrabure y Correa (Eds.), Coleccion de Leyes, Decretos y Besoluciones Lima, Peru,14, 328-351.

Rengifo-Salgado, E., Rios-Torres, S., Fachín Malaverri, L., & Vargas-Arana, G. (2017). Saberes ancestrales sobre el uso de flora y fauna en la comunidad indígena Tikuna de Cushillo Cocha, zona fronteriza Perú-Colombia-Brasil. Revista peruana de biología, 24(1), 67-78.

Renner, S.S., H. Balslev, and L.B. Holm-Nielson. (1990). Flowering Plants of Amazonian Ecuador: A Checklist. AAU Reports #24, Botanical Institute, University of Aarhus, Denmark.

Revilla, J. (2002) Plantas uteis da bacia amazonica. Manaus. SEBRAE, INPA

Reyes-Garcia, V. E. (2001). Indigenous people, ethnobotanical knowledge, and market economy: a case study of the Tsimane Amerindians in lowland Bolivia. University of Florida.

Reyes, C. (2017). Conocimiento y uso de plantas en tres comunidades Kichwas: Yana Yaku, Loro Cachi y Nina Amarun, Pastaza-Ecuador. Cinchonia, 15(1), 164-256.

Ribeiro de Sampaio, F. X., (1825) Diario da Viagem, que em visita, e correicao das povoacoes da capitania de S. Jose do Rio Negro, 1774 1775, Typografia de Academia, Lisboa, p. 34.

Ribeiro Magno-Silva, E., Teixeira Rocha, T., & Caldeira Tavares-Martins, A. C. (2020). Ethnobotany and ethnopharmacology of medicinal plants used in communities of the soure marine extractive reserve, Pará State, Brazil. Boletín Latinoamericano y del Caribe de Plantas Medicinales y Aromáticas, 19(1).

Ribeiro, R. V., Bieski, I. G. C., Balogun, S. O., & de Oliveira Martins, D. T. (2017). Ethnobotanical study of medicinal plants used by Ribeirinhos in the North Araguaia microregion, Mato Grosso, Brazil. Journal of ethnopharmacology, 205, 69-102.

Rios, M., and J. Caballero (1997) Las plantas en la alimentación de la comunidad Ahuano, Amazonía ecuatoriana. Uso y manejo de recursos vegetales-Memorias del segundo simposio ecuatoriano de etnobotánica y botánica económica. Ediciones Abya-Yala, Quito.

Ritter, R.A. et al. (2012) Ethnoveterinary knowledge and practices at Colares island, Pará state, eastern Amazon, Brazil." Journal of Ethnopharmacology 144: 346-352.

Rizzini, G.T. & W. Mors (1976) Botànica Econòmica Brasileira São Paulo, EPU, Ed. da Universidade do São Paulo. Pages 96-99

Robineau, L. Ed. (1991) Towards a Caribbean Pharmacopoeia. TRAMIL-4 WORKSHOP, UNAH, Enda Caribe, Santo Domingo

Rodrigues de Freitas, R., Haverroth, M. & Siviero, A. (2016). Educação agroflorestal para resiliência socioecológica de Reservas Extrativistas da Amazônia. Chapter 3.In: da Silva Ramos, J. I., Cavalcante, O. P., da Silva, J. O., de Oliveira, M. A., Fernandes, C. C., da Silva, J. P., & Uchôa, J. M. S. (2016). Etnobotânica e botânica econômica do Acre ISBN: 978-85-8236-027-9 Copyright© Edufac 2016, Amauri Siviero, Lin Chau Ming, Marcos Silveira, Douglas Daly, Richard Wallace (Organizadores) Editora da Universidade Federal do Acre-Edufac.

Rodrigues, E. (2006) Plants and animals utilized as medicines in the Jaú National Park (JNP), Brazilian Amazon. Phytotherapy Research: An International Journal Devoted to Pharmacological and Toxicological Evaluation of Natural Product Derivatives, 20(5), pp.378-391.

Rodrigues, E., Duarte-Almeida, J. M., & Pires, J. M. (2010). Perfil farmacológico e fitoquímico de plantas indicadas pelos caboclos do Parque Nacional do Jaú (AM) como potenciais analgésicas: parte I. Revista Brasileira de Farmacognosia, 20, 981-991.

Rodrigues, E., Mendes, F. R., & Negri, G. (2006). Plants indicated by brazilian indians for disturbances of the central nervous system: A bibliographical survey. Central Nervous System Agents in Medicinal Chemistry (Formerly Current Medicinal Chemistry-Central Nervous System Agents), 6(3), 211-244.

Rodrigues, I., & Oliveira, A. E. D. (1977). Alguns aspectos da ergologia Mura-Pirahã. Bol. Museu Paraense Emilio Goeldi 65: 1-54.

Rodrigues, S., Caetano N, D. G., & Caetano, C. M. (2007). Espécies frutíferas do centro-sul do Estado de Rondônia, Amazônia brasileira. Acta Agronómica, 56(2), 69-74.

Rodriquez, F. (1996) Waorani hunting and harvesting practices in Ecuador. CTFS/STRI

Rojas, R., G. Ruiz , P. Ramírez, C. F. Salazar, C. Rengifo, C. Ll. Flores, C. Marín, D. Torres, J. Ojanama, W. Silvano, V. Muñoz, H. Luque , N. Vela, N. del Castillo, J. Solignac, V. R. López de Oliveira,and F. de María Panduro (2001) Comercialización De Masa Y √áFruto Verde√à De Aguaje (Mauritia flexuosa L.F.) En Iquitos (Perú). Folia Amazonica 12: 15-38

Román, F.J. (2002) Especies forestales utilizadas en la construcción de la vivienda tradicional asháninka en el ámbito del Río Perené (Junín, Perú). Facultad de Ciencias Forestales. Universidad Nacional Agraria La Molina, Lima.

Romanoff, S., D. Manquid, F. Shoque, and D.W. Fleck (2004) La vida tradicional de los Matsés. CAAAP, Lima

Roth, W.E. (1924). An introductory study of the arts, crafts and customs of the Guiana Indians. 38th Ann. Rep. Bur. Am. Ethnol. 1916-17: 25-745.

Ruiz, L., Ruiz, L., Maco, M., Cobos, M., Gutierrez-Choquevilca, A. L., & Roumy, V. (2011). Plants used by native Amazonian groups from the Nanay River (Peru) for the treatment of malaria. Journal of Ethnopharmacology, 133(2), 917-921.

Rusby, H.H. (1924). Miré. Journal of the American Pharmaceutical Association 13: 101-102

Rutter, R.A. (1990). Catalogo de plantas utiles de la Amazonia peruana. Comunidades y Culturas Peruanas No. 22. Ministerio de Edudación, Instituo Linguistico de Verano.

Sánchez, M. (1997). Catálogo preliminar comentado de la flora del Medio Caquetá. Estudios en la Amazonia colombiana, 12.

Sánchez, M. (2005) Use of tropical rain forest biodiversity by indigenous communities in northwestern Amazonia. Universiteit van Amsterdam/COLCIENCIAS, Bogotá

Sánchez, M., and P. Miraña (1991) Utilización de la vegetación arbórea en el Medio Caquetá: 1. El árbol dentro de las unidades de la tierra, un recurso para la comunidad Miraña. Colombia Amazonica 5: 69-98

Saltos, R. V. A., Vásquez, T. E. R., Lazo, J. A., Banguera, D. V., Guayasamín, P. D. R., Vargas, J. K. A., & Peñas, I. V. (2016). The use of medicinal plants by rural populations of the Pastaza province in the Ecuadorian Amazon. Acta Amazonica, 46, 355-366.

Sampaio, A. J. de. (1916). A flora de Matto Grosso. Archiv. Mus. Nac. Rio de Janeiro 19:

San Sebastián, M. (1995) Ñucanchic Janpi- Tratamientos con plantas medicinales de los Naporunas. CICAME- SANDI YURA, Coca

Santos, M. R. A., Lima, M. R., & Oliveira, C. L. L. G. (2014). Medicinal plants used in rondônia, Western amazon, Brazil. Revista Brasileira de Plantas Medicinais, 16(3), 707-720. Chicago

Santos, M., Pantoja, T. R., & Oliveira, A. M. (2022). Uso Popular De Plantas Medicinais No Tratamento Do Diabetes Mellitus No Estado Do Amapá, Brasil. Resumos XXVI SPMB.

Sanz-Biset, J. et al. (2009) A first survey on the medicinal plants of the Chazuta valley (Peruvian Amazon). Journal of Ethnopharmacology 122.2: 333-362.

Schultes, R. E. (1977). Diversas plantas comestíveis nativas do noroeste da Amazônia. Acta amazônica, 7, 317-327.

Schultes, R. E., & Raffauf, R. F. (1992). A rare report of an intoxicating snuff from the Amazon. Kew Bulletin 47(4): 743-744

Schultes, R.E. (1951) Plantae Austro-Americanae VII. Botanical Museum Leaflets 15: 29-78

Schultes, R.E. (1955). A new generic concept in the Euphorbiaccae. Bot. Mus. Leafl., Harvard Univ., 17: 27√°36.

Schultes, R.E. (1956). The Amazon Indian and evolution in Hevea and related genera. Journ. Arn. Arb .37: 123-152.

Schultes, R.E. (1974) Palms and religion in the Northwest Amazon. Principes 18: 3-21

Schultes, R.E. (1976) Planlae Colombianae XIX. E partibus amazonicis witotorum plantae fructuariae sativae novae". Bot. Mus. Leafl. Harvard Univ. 24: 193-204.

Schultes, R.E. & Raffauf, R. F. (1990). Field notes on curare constituents in the Northwest Amazonia. Curare, 13(2), 105-120

Schultes, R.E. & Raffauf, R.F. (1990). The Healing Forest: Medicinal and Toxic Plants of the Northwest Amazonia (Dioscorides Press, Portland, OR).

Schwacke, W. (1884). Bereitung des Curare-Pfeilgiftes bet den Tecuna-Indianern. Jahrb. Bot. Gart. Berlin 3: 220-223.

Seoane, E., and S. Soplín (1999) Plantas medicinales utilizadas en la regulación de la fertilidad. Biota 17: 82-99

Shepard, G.H., D.W. Yu, M. Lizarralde, et al. (2001) Rain forest habitat classification among the Matsigenka of the Peruvian Amazon. Journal of Ethnobiology 21: 1-38

Shepard, G.H., M.N.F. da Silva, A.F. Brazao, and P. van der Veld. (2004). Arte Baniwa: Sustentabilidade socioambiental de aruma no Alto Rio Negro. In Terras ind√µgenas e unidades de conservacao danatureza:O desafio das sobreposicoes, ed. F. Ricardo, pp. 129-143. Instituto Socioambiental, Sao Paulo.

Shrestha, T., Kopp, B., & Bisset, N. G. (1992). The Moraceae-based dart poisons of South America. Cardiac glycosides of Maquira and Naucleopsis species. Journal of ethnopharmacology, 37(2), 129-143.

Silva Farias, M. de Oliveira, L.C., de Melo Figueiredo, S.M., Rodrigues Pereira, L. & Rodrigues, E (2016) Diversidade e uso de palmeiras da mata ciliar do rio Acre. Chapter 7. In: da Silva Ramos, J. I., Cavalcante, O. P., da Silva, J. O., de Oliveira, M. A., Fernandes, C. C., da Silva, J. P., & Uchôa, J. M. S. (2016). Etnobotânica e botânica econômica do Acre ISBN: 978-85-8236-027-9 Copyright© Edufac 2016, Amauri Siviero, Lin Chau Ming, Marcos Silveira, Douglas Daly, Richard Wallace (Organizadores) Editora da Universidade Federal do Acre-Edufac.

Silva, H., and J. García (1997) La Medicina Tradicional en Loreto. Instituto Peruano de Seguridad Social / Instituto de Medicina Tradicional, Iquitos

Silva, R. B. L. (2010). Diversidade, uso e manejo de quintais agroflorestais no Distrito do Carvão, Mazagão-AP, Brasil. 284 f. Tese (Doutorado em Desenvolvimento Sustentável do Trópico √≤mido) - Universidade Federal do Pará/Núcleo de Altos Estudos Amazônicos, Belém.

Silva, R. B. L. (2002). A etnobotânica de plantas medicinais da comunidade quilombola de Curiaú, Macapá-AP, Brasil. 170 f. Dissertação (Mestrado em Agronomia) - Departamento de Biologia Vegetal, Faculdade de Ciências Agrárias do Pará, Belém.

Silva, R. B. L. et al. (2013). Caracterização agroecológica e socioeconômica dos moradores da comunidade quilombola do Curiaú, Macapá-AP, Brasil. Biota Amazônia, Macapá, v. 3, n. 3, p. 113-138.

Silva, W. et al. (2019) Guia etnobotânico de plantas em comunidades Desano (Tukano-oriental) no rio Tiquié - Brasil. In: Cadernos de Etnolingüística, vol. 7, n. 1, p. 1-42

Skov, F., and H. Balslev (1989) A revision of Hyospathe (Arecaceae). Nordic Journal of Botany 9: 189-202

Smith, A. C. (1939). Notes on a collection of plants from British Guiana. Lloydia, 2: 161-218.

Smith, N. (1979). A Pesca no Rio Amazonas. INPA, Manaus. Pg. 49

Smith, N., R. Vásquez, and W. H. Wust (2007) Amazon river fruits. Flavors for Conservation. Amazon Conservation Association (ACA)/ Missouri Botanical Garden Press, Lima

Soares Machado, F. (2016) Etnobotânica de espécies florestais não madeireiras em comunidades locais do Vale do Juruá, Acre. Chapter 2. In: da Silva Ramos, J. I., Cavalcante, O. P., da Silva, J. O., de Oliveira, M. A., Fernandes, C. C., da Silva, J. P., & Uchôa, J. M. S. (2016). Etnobotânica e botânica econômica do Acre ISBN: 978-85-8236-027-9 Copyright© Edufac 2016, Amauri Siviero, Lin Chau Ming, Marcos Silveira, Douglas Daly, Richard Wallace (Organizadores) Editora da Universidade Federal do Acre-Edufac.

Sosnowska, J, D. Ramírez & B. Millán. (2010). Palmeras usadas por los indígenas Asháninkas en la Amazonía Peruana. Revista Peruana de Biología 17(3): 347-352.

Soukup, J. (1970) Vocabulary of the Common Names of the Peruvian Flora and Catalog of the Genera. Editorial Salesiano, Lima. 436 pp.

Souto, R. N. P. S. et al. (2011). Estudos preliminares da atividade inseticida de óleos essenciais de espécies de Piper linneus (piperaceae) em operárias de Solenopis saevissima f Smith (Hymenoptera: formicidae ), em laboratório. Biota Amazônica. Macapá, v. 1, n. 1, p. 42-48.

Souza, B. (1956). O cipó-babão (Cissus gongylodes Baker). Um agente coagulante do látex de Hevea. Bol. Técnico do Instituto Agronômico do Norte (31): 163-186.

Spruce, R. (1853) Notes of a Botanist on the Amazon and Andes, Macmillan, London, 1908.

Stagegaard, J., M. Sørensen, and L.P. Kvist (2002) Estimations of the importance of plant resources extracted by inhabitants of the Peruvian Amazon flood plains. Perspectives in Plant Ecology, Evolution and Systematics 5: 103-122

Steward, JH and Metraux, A (1948) Tribes of the Peruvian and Ecuadorian Montaña. In HJ Steward, ed. Handbook of South American Indians. Vol 3: The Tropical Forest Tribes. Bureau of American Ethnology. Bulletin 143, Washington DC. Pp. 594, 605.

Steyemark, J.A. (1984) Flora de Venezuela. Piperaceae. Ediciones Fundacion Ambiental, Venezuela

Svenning, J.C., and M.J. Macía (2002) Harvesting of Geonoma macrostachys Mart. leaves for thatch: an exploration of sustainability. Forest Ecology and management 167: 251-262

Thevet, A. (1557). Les singularites de la France antarctique, autrement nommee Amerique; et Isles decouvertes de nostre temps. Nouvelle edition avec notes et commentaires par Paul Gaffarel. Paris. (Paris ed. 1557; Anvers ed. 1558)

Thomas, E. , and I. Vandebroek (2006) Guía de Plantas Medicinales de los Yuracarés y Trinitarios del Territorio Indígena Parque Nacional Isiboro-Sécure, Bolivia. Industrias gráficas Sirena, Santa Cruz

Thomas, E. (2008) Quantitative Ethnobotanical Research on Knowledge and Use of Plants for Livelihood among Quechua, Yuracaré and Trinitario Communities in the Andes and Amazon Regions of Bolivia.. Faculty of Bioscience Engineering. Ghent University, Belgium

Ticona, J. P. (2001) Los chimane: conocimiento y uso de plantas medicinales en la comunidad Tacuaral del Matos ( Provincia Ballivián, Departamento del Beni). Facultad de Ciencias Puras y Naturales. Universidad Mayor de San Andres, La Paz

Tomchinsky, B. (2014). Etnobotânica de plantas antimaláricas em Barcelos, Amazonas.

Tomchinsky, B., Ming, L. C., Kinupp, V. F., Hidalgo, A. D. F., & Chaves, F. C. M. (2017). Ethnobotanical study of antimalarial plants in the middle region of the Negro River, Amazonas, Brazil. Acta Amazonica, 47(3), 203-212

Tournon, J. (2006) Las Plantas, los Rao y sus espíritus ( Etnobotánica del Ucayali). Gobierno Regional de Ucayali, Pucallpa

Tournon, J., Raynal-Roques, A., & Zambettakis, C. (1986). Les Cyperacees medicinales et magiques de L'Ucayali. Journal d'agriculture traditionnelle et de botanique appliquée, 33(1), 213-224.

Townsend, W.R. (1996) Nyao Itõ: caza y pesca de los Sirionó. Instituto de Ecología, Universidad Mayor de San Andrés, La Paz

Triana, G. (1985) Los Puinaves del Inirida. Formas de subsistencia y mecanismos de adaptación.. Instituto de Ciencias Naturales-Museo de Historia Natural. Universidad Nacional de Colombia, Bogotá.

Trujillo, W. & Correa-Munera, M. (2010). Plants used by a Coreguaje indigenous community in the Colombian Amazon. Caldasia, 32(1), 1-20.

Tudela-Talavera, P., & La Torre, M. D. L. A. (2015). Cultural importance and use of medicinal plants in the Shipibo-Conibo native community of Vencedor (Loreto) Peru. Ethnobotany Research and Applications, 14, 533-548.

Vásquez, M., and J. B. Vásquez (1998) La extraccíon de productos forestales diferentes de la madera en el ambito de Iquitos-Perú. Folia Amazonica 9: 69-84

Vásquez, R. (1992). Sistemática de las plantas medicinales de uso frecuente en el área de Iquitos. Folia Amazónica 4: 65-80.

Vásquez, S. P. F., Mendonça, M. S. D., & Noda, S. D. N. (2014). Etnobotânica de plantas medicinais em comunidades ribeirinhas do Município de Manacapuru, Amazonas, Brasil. Acta amazônica, 44, 457-472.

Vázquez, M.R. (1990) Useful Plants of Amazonian Peru. Spanish Typescript. Second Draft. Filed with USDA's National Agricultural Library.

Vélez, G.A., and A.J. Vélez (1999) Sistema agroforestal de las chagras indígenas del Medio Caquetá. TROPENBOS

Valadeau, C., J. Alban Castillo, M. Sauvainc, A. Francis Lorese & G. Bourdy. (2010). The rainbow hurts my skin: Medicinal concepts and plants uses among the Yanesha (Amuesha), an Amazonian Peruvian ethnic group. Journal of Ethnopharmacology 127: 175-192.

Valadeau, C., Pabon, A., Deharo Eric, Albán-Castillo, J., Estevez, Y., Lores, F.A., Rojas, R., Gamboa, D., Sauvain, M., Castillo, D., Bourdy, G. (2009). Medicinal plants from the Yanesha (Peru): Evaluation of the leishmanicidal and antimalarial activity of selected extracts. Journal of Ethnopharmacology 123, 413-422.

Valero, H. (1969). Yanoáma: the story of a woman abducted by Brazilian Indians. Allen & Unwin, London.

Valle, J.R.& N.P.Silva (1973). Ichthyotoxicity of cannabinoids. Cienc. e Cult. 25: 647.

Van Andel, T. R. (2000). Non-timber forest products of the North-West District of Guyana. Utrecht University.

van den Berg, M. E. (1984). Ver-o-peso: the ethnobotany of an Amazonian market. Advances in economic botany, 1, 140-149.

van den Berg, M. E. (1993). Plantas medicinais na Amazônia(contribuição ao seu conhecimento sistemático). Coleção Adolpho Ducke. Belém, Museu Paranaense Emilio Goeldi. 207 pg. ISBN: 85-7098-041-8

van den Berg, M. E., & Silva, M. H. L. D. (1988). Contribuição ao conhecimento da flora medicinal de Roraima. Acta amazônica, 18, 23-35.

Van den Eynden, V., E. Cueva, and O. Cabrera (2004) Edible palms of Southern Ecuador. Palms 48: 141-157

Van der Linden, M., and R. López (1990) Utilización de palmeras amazónicas en el nororiente peruano.. Revista Forestal del Perú 17: 65-74

Vargas, G. (2006) Transformación Y Elaboración De Alimentos Con Especies Vegetales Y Animales Por Las Comunidades Cubeas Del Cuduyari. Instituto amazónico de investigaciones científicas-Sinchi, Bogotá.

Vargas, L. (1997) Vida y medicina tradicional de los Mosetenes de Muchanes. Ecología en Bolivia 29: 19-44

Vasquez and Gentry, cited in Duke's Amazonian Ethnobotanical Dictionary

Vasquez, R., and A.H. Gentry (1989) Use and misuse of forest-harvested fruits in the Iquitos area. Conservation Biology 3: 350-361

Veiga, J. B. D. (2011). Etnobotânica e etnomedicina na Reserva de Desenvolvimento Sustentável do Tupé, baixo rio Negro: plantas antimaláricas, conhecimentos e percepções associadas ao uso e à doença.

Vickers, W.T., and T. Plowman (1984) Useful plants of the Siona and Secoya indians of Eastern Ecuador. Fieldiana, Botany 15: 1-63

Villa Muñoz, G., Garwood, N.C, Bass, M.S., & Navarrete Zambrano, H. G. (2016). Common trees of Yasuní. A guide for identifying the common trees of the Ecuadorian Amazon.

von Martius, C.F.P. (1843) Beitrage zur Kenntmss der Gattung Erythroxyton. Abhandlungen der mathematisch-physikalischen Classe der koniglich bayerischen Akademie der Wissenschaften, Munich, 3: 367 - 369.

Von Reis, S. (1973) Drugs & foods from little-known plants. Notes in Harvard University Herbaria. Harvard University Press, Cambridge, Massachusetts.

Von Reis, S. & F.J. Lipp (1982) New plant sources for drugs and foods from the New York Botanical Garden Herbarium. Harvard University Press. Cambridge. Massachusetts.

Vormisto, J. (2002) Making and marketing chambira hammocks and bags in the village of Brillo Nuevo, northeastern Peru. Economic Botany 56: 27-40

Wallace, A.R. (1853). A Narrative of Travels on the Amazon and Rio Negro. London, Reeve and Co.

Webster, L.J. (1970) Letter to R.E. Schultes. February 5, 1970

Wheeler, M.A. (1970) Siona use of chambira palm fiber. Economic Botany 24: 180-181

Williams, E.P. (1962). Algunos datos sobre el Barbasco. Bol. Soc. Venez. Cienc. Nat. 6: 21- 34.

Wurdack, J. J. (1958). Indian narcotics in southern Venezuela. Gard. Journ., 116-118.

Xavier, W. K. S.; Cunha, E. D. S. (2015). Comercialização de produtos naturais medicinais oriundos do Estado do Amapá. Biota Amazônica, Macapá, v. 5, n. 2, p. 23-25.

Yanomami, M.I., Yanomami, E., Albert, B., Milliken, W., & Coelho, V. (2015). Hwër_mamotima thë pë ã oni= Manual dos remédios tradicionais Yanomami. São Paulo: Boa Vista, Brazil

Zambrana, N. Y. P., Zambrana, N. Y. P., & Bussmann, R. W. (2018). La Etnobotánica de los Chácobo en el Siglo XXI. Ethnobot. Res. Appl, 16, 1-149.

Zarucchi, J. L. (1980). Ibapichuna: an edible Dacryodes (Burseraceae) from the northwest Amazon. Botanical Museum Leaflets, Harvard University, 28(1), 81-85.

Zent, E. M. L. (1999). Hoti ethnobotany: Exploring the interactions between plants and people in the Venezuelan Amazon (Doctoral dissertation, University of Georgia)
